# NLRP3 activation in response to disrupted endocytic traffic

**DOI:** 10.1101/2021.09.15.460426

**Authors:** Bali Lee, Christopher Hoyle, Jack P. Green, Rose Wellens, Fatima Martin-Sanchez, Daniel Williams, Paula I. Seoane, Hayley Bennett, Antony Adamson, Gloria Lopez-Castejon, Martin Lowe, David Brough

**Affiliations:** Division of Neuroscience and Experimental Psychology, School of Biological Sciences, Faculty of Biology, Medicine and Health, University of Manchester, Manchester Academic Health Science Centre, Manchester, UK; Division of Infection, Immunity and Respiratory Medicine, School of Biological Sciences, Faculty of Biology, Medicine and Health, Manchester Academic Health Science Centre, University of Manchester, Manchester, UK; Division of Molecular and Cellular Function, School of Biological Sciences, Faculty of Biology, Medicine and Health, Manchester Academic Health Science Centre, University of Manchester, Manchester, UK; Geoffrey Jefferson Brain Research Centre, The Manchester Academic Health Science Centre, Northern Care Alliance NHS Group, University of Manchester, Manchester, UK; The Lydia Becker Institute of Immunology and Inflammation, University of Manchester, Manchester, UK; Department of Biomedical Science, Centre for Membrane Interactions and Dynamics, University of Sheffield, Firth Court, Sheffield S10 2TN, UK; Genome Editing Unit, Faculty of Biology, Medicine and Health, University of Manchester, Manchester, UK

## Abstract

Inflammation driven by the NLRP3 inflammasome is coordinated through multiple signaling pathways and with a poorly defined regulation by sub-cellular organelles. Here, we tested the hypothesis that NLRP3 senses disrupted endosome trafficking to trigger inflammasome formation and inflammatory cytokine secretion. NLRP3-activating stimuli disrupted endosome trafficking and triggered localization of NLRP3 to vesicles positive for endosome markers and the inositol lipid PtdIns4P. Chemical disruption of endosome trafficking sensitized macrophages to the NLRP3 activator imiquimod driving enhanced inflammasome activation and cytokine secretion. Together these data suggest that NLRP3 is capable of sensing disruptions in the trafficking of endosomal cargoes, and that this may explain in part the spatial activation of the NLRP3 inflammasome complex. These data highlight new mechanisms amenable for the therapeutic targeting of NLRP3.

**One-Sentence Summary:** NLRP3 senses disruptions in endosome trafficking to trigger the formation of an inflammasome complex and initiate an inflammatory response.

## Main Text

The NLRP3 (NLR family pyrin domain containing 3) inflammasome is a multi-molecular protein complex that regulates the processing and secretion of the pro-inflammatory cytokines interleukin (IL)-1β and IL-18. NLRP3 inflammasome activation is commonly reported in cells of the innate immune system, where the cytosolic protein NLRP3 senses diverse cellular stressors and undergoes post-translational modifications and a conformational change to enable interaction with an adaptor protein called ASC (Apoptosis-associated speck-like protein containing a CARD). The subsequent oligomerisation of ASC into a ‘speck’ provides a platform for the recruitment and activation of the protease caspase-1, which cleaves inactive precursors of IL-1β and IL-18 (pro-forms) to facilitate their release from the cell (e.g. (*1*)). NLRP3-dependent inflammation contributes to the worsening of diverse diseases and although there are inhibitors of NLRP3 being developed (*2*) we do not fully understand the mechanisms leading to its activation.

In macrophages, activation of the canonical NLRP3 inflammasome requires two stimuli. The first is required to induce expression of the NLRP3 protein itself in a step known as priming, and experimentally this is usually achieved by treatment of cells with the pathogen-associated molecular pattern (PAMP) bacterial endotoxin (or lipopolysaccharide, LPS) (*3*). For an active NLRP3 inflammasome to assemble the primed cell needs to then encounter a further PAMP, or damage-associated molecular pattern (DAMP), or another activating stimulus. Structurally diverse PAMPs, DAMPs, and other activators are reported to activate NLRP3, with a suggested common consequence of their action an alteration in cellular homeostasis that is sensed by NLRP3 (*4*). We recently reviewed literature highlighting that multiple organelle stresses are linked to NLRP3 inflammasome activation, and suggested a potentially under-appreciated importance of endosomes in the process (*1*). Dispersion of the *trans-Golgi* marker TGN38 (in primates this is TGN46) occurs after treatment of cells with NLRP3-activating stimuli (*5*). This is suggested to represent a dispersed *trans*-Golgi network (TGN) that is important for NLRP3 inflammasome activation (*5*). However, TGN38/46 does not permanently reside in the TGN, and cycles through the plasma membrane and endosomes before returning to the TGN. Cycling of TGN38/46 is interrupted by modification of endosomal pH using vacuolar ATPase inhibitors, or K^+^ ionophores, both of which are known to cause an accumulation of TGN38/46 in early endosomes (*6, 7*). K^+^ ionophores are also very well established activators of NLRP3 (*8*), and other activators of NLRP3 (such as hypo-osmotic stress, and the K^+^-efflux-independent stimuli imiquimod and CL097), also perturb endosomal pH (*9, 10*). Thus, inflammasome activation may also be associated with perturbed endosome function. There is also extensive evidence that endosomes provide a signalling hub for pattern recognition receptors (PRRs) such as members of the Toll-Like Receptor (TLR) family, identifying this organelle as a site of importance for inflammatory signalling (*11*). We therefore tested the hypothesis that NLRP3 senses disrupted endosome trafficking as a trigger for its activation.

## Results

### TGN38/46 trapping in endosomes

Initially we set out to test the hypothesis that NLRP3-activating stimuli cause trapping of TGN38/46 in the endosome. To do this we first assembled a panel of diverse NLRP3 activating stimuli. Included as NLRP3 activating stimuli were the K^+^ ionophore nigericin (Nig) (*12*), lysosomal disrupting agent L-leucyl-L-leucine methyl ester (LLOMe) (*13*), and imiquimod (IQ), which activates NLRP3 independently of K^+^ efflux (*10*). The Na^+^ ionophore monensin is known to disrupt endosome trafficking (*7*), and as such was included as a control. COS7 cells were treated with the panel of stimuli for 90 minutes, and then labelled with antibodies for TGN46, and the permanent TGN resident protein Golgin-97 (*14*). In unstimulated cells, there was strong co-localization between Golgin-97 and TGN46, with both proteins localizing to the peri-nuclear TGN, as expected (Fig. 1A&B). Treatment with NLRP3-activating stimuli or monensin caused dispersal of TGN46 to cytoplasmic puncta, and a reduction in co-localization with Golgin-97, which remained peri-nuclear (Fig. 1A&B, fig. S1A). Imiquimod had the mildest effect of the NLRP3-activating stimuli tested on TGN46 localization (Fig. 1B, fig. S1A). These data suggest that under these conditions, in response to NLRP3-activating stimuli, the TGN, as defined by Golgin-97, remains largely intact, but that TGN46 becomes vesicular. As discussed, TGN46 becomes trapped in early endosomes when endosome to Golgi trafficking is perturbed (*6, 7*). Thus, we investigated whether the TGN46 puncta represented endosomes, using the early endosomal marker EEA1. In untreated cells there was little co-localization between TGN46 and EEA1 (Fig. 1C&D). However, in response to the NLRP3-activating stimuli or monensin, there was a significant increase in TGN46 and EEA1 co-localization (Fig. 1C&D, fig. S1B). The NLRP3-activating stimuli and monensin also affected the distribution of CD63-positive late endosomes, which became more scattered after stimulation (Fig. 1E, fig. S1C). These scattered CD63 endosomes also co-localized with TGN46 (Fig. 1E, fig. S1C), although this increase in co-localization was not significant as assessed by quantitation due to coincidental clustering of the CD63 endosomes around the peri-nuclear TGN prior to stimulation (Fig. 1F). A similar effect to that observed with CD63 was observed when we examined TGN46 co-localization with the lysosomal marker LAMP1, with the exception of LLOMe, likely due to its role as a lysosomal disrupting agent (fig. S2A&B). There was no increase in co-localization with the recycling endosome marker MICAL-L1 (fig. S2C&D). These data indicate that trapping of TGN46 in endocytic compartments is a common outcome of treatment with NLRP3-activating stimuli.

**Fig. 1.**
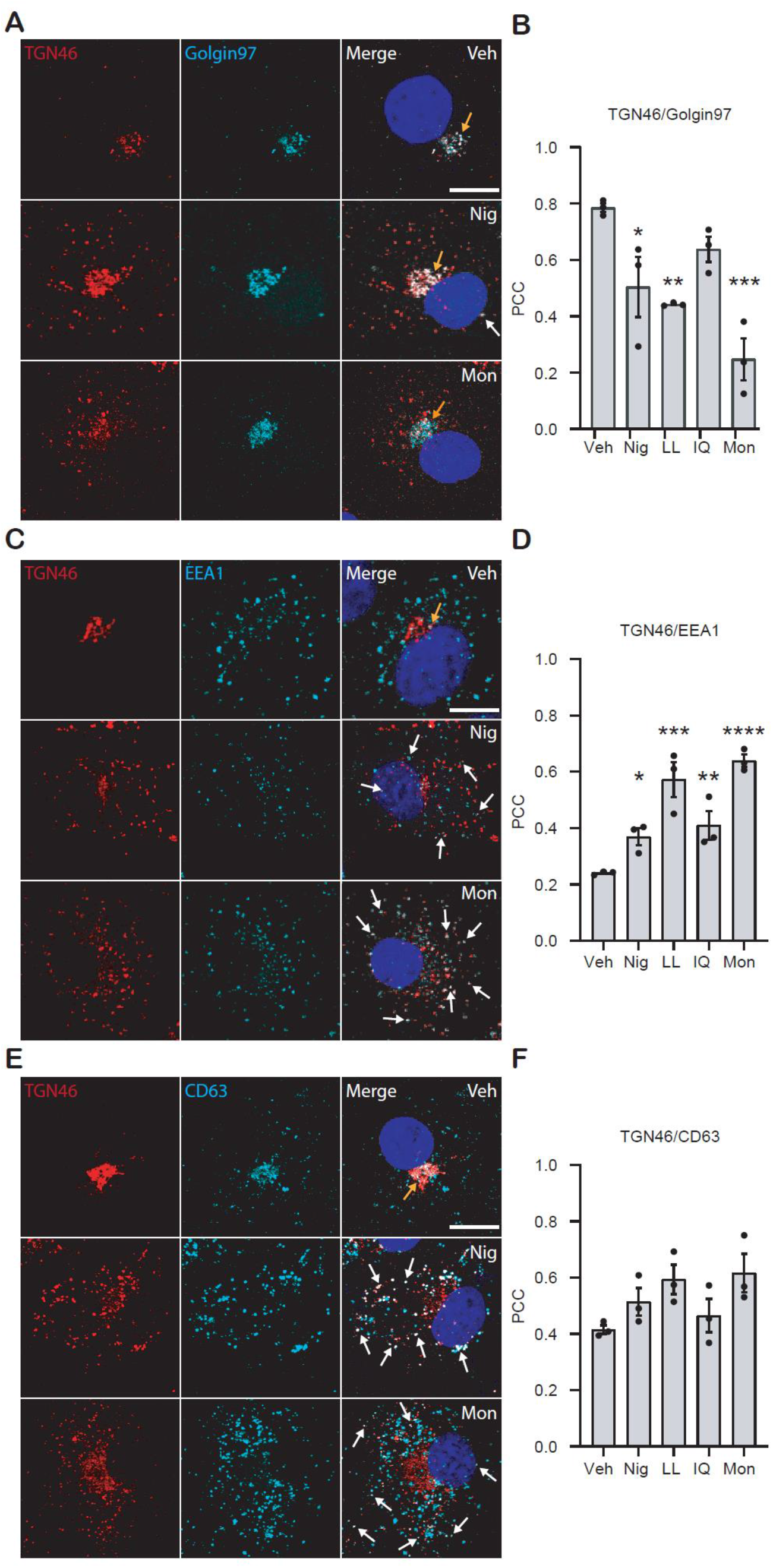
TGN46 co-localizes with endolysosomal membrane markers after stimulation of COS7 cells with NLRP3-activating stimuli. COS7 cells were stimulated for 90 min with vehicle (Veh, 0.5% ethanol (v/v)), nigericin (Nig, 10 μM), LLOMe (LL, 1 mM), imiquimod (IQ, 75 μM) or monensin (Mon, 10 μM). **(A)** Immunofluorescence images following vehicle, nigericin or monensin treatment showing the co-localization of TGN46 with Golgin-97. See also fig. S1. **(B)** Pearson’s correlation coefficient (PCC) between TGN46 and Golgin-97 for all stimuli. **(C)** Immunofluorescence images following vehicle, nigericin or monensin treatment showing the co-localization of TGN46 with EEA1. See also fig. S1. **(D)** Pearson’s correlation coefficient (PCC) between TGN46 and EEA1 for all stimuli. **(E)** Immunofluorescence images following vehicle, nigericin or monensin treatment showing the co-localization of TGN46 with CD63. See also fig. S1. **(F)** Pearson’s correlation coefficient (PCC) between TGN46 and CD63 for all stimuli. All images are representative of 3-5 independent experiments. Blue represents nuclei staining by DAPI, yellow arrowheads indicates co-localization at the Golgi, white arrowheads indicate co-localization on puncta. Scale bars: 10 μm. Values are mean ± SEM (n = 3). One-way ANOVA followed by Bonferroni’s multiple comparison test was performed for data analysis. *: P≤ 0.05, **: P≤ 0.01, ***: P≤0.001 and ****: P≤0.0001.

We next assessed the effects of the stimuli on TGN38 distribution in primary mouse bone marrow-derived macrophages (BMDMs), cells commonly used to study NLRP3 inflammasome activation. In untreated and LPS-treated BMDMs, Golgin-97 and TGN38 co-localized in the peri-nuclear TGN (Fig. 2A&B). Treatment with NLRP3-activating stimuli (except imiquimod) or monensin caused a decrease in Golgin-97 and TGN38 co-localization, with a concomitant increase in TGN38 dispersion, similar to that seen in COS7 cells (Fig. 2A&B, fig. S3A). Treatment with NLRP3 activating stimuli did not cause quantitatively significant co-localization of TGN38 with EEA1, though some co-localization was detected (Fig. 2C&D, fig. S3B). In LPS-treated BMDMs, nigericin, LLOMe, and imiquimod all induced significant IL-1β release and this was inhibited by the NLRP3 inhibitor MCC950 (Fig. 2E). Monensin did not induce IL-1β release despite inducing dispersal of TGN38, suggesting that TGN38 dispersal alone is not sufficient for NLRP3 activation.

**Fig. 2.**
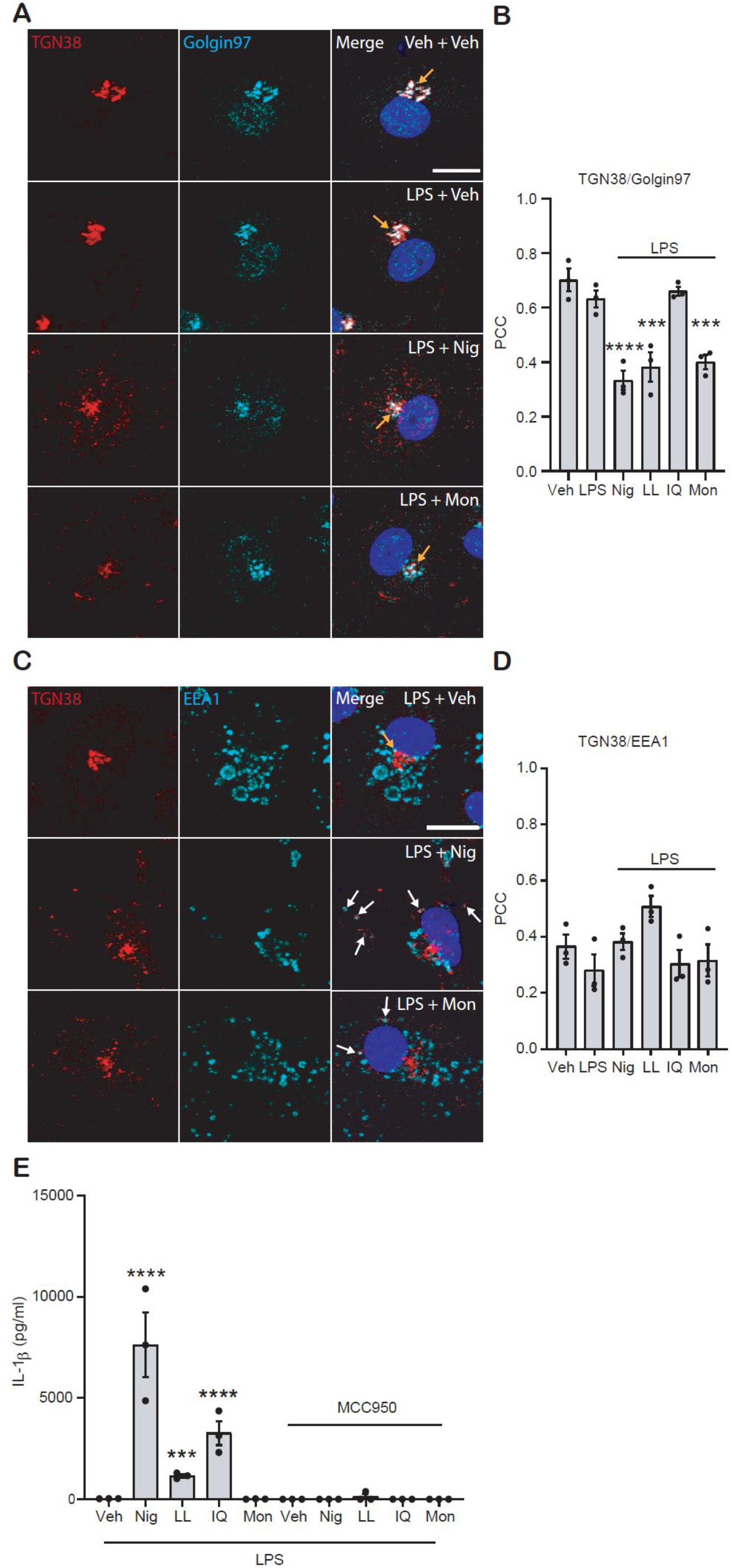
TGN38 co-localizes with EEA1 after stimulation of BMDMs with NLRP3-activating stimuli. BMDMs were primed with LPS (1 μg mL^−1^), with the exception of vehicle, for 4 h. In order to prevent NLRP3-dependent cell death to aid the collection of immunofluorescence images, MCC950 (10 μM) was applied 15 min prior to a 90 min stimulation with vehicle (Veh, 0.5% ethanol (v/v)), nigericin (Nig, 10 μM), LLOMe (LL, 1 mM), imiquimod (IQ, 75 μM) or monensin (Mon, 10 μM). **(A)** Immunofluorescence images following vehicle, nigericin or monensin treatment showing the co-localization of TGN38 with Golgin-97. See also fig. S3. **(B)** Pearson’s correlation coefficient (PCC) between TGN38 and Golgin-97 for all stimuli. **(C)** Immunofluorescence images following vehicle, nigericin or monensin treatment showing the co-localization of TGN38 with EEA1. See also fig. S3. **(D)** Pearson’s correlation coefficient (PCC) between TGN38 and EEA1 for all stimuli. **(E)** IL-1β release in response to stimuli. All images are representative of 3-5 independent experiments. Blue represents nuclei staining by DAPI, yellow arrowheads indicate instances of co-localization at the Golgi, white arrowheads indicate co-localization on puncta. Scale bars: 10 μm. IL-1β release was measured by ELISA. Values are mean ± SEM (n = 3). One-way ANOVA followed by Bonferroni’s multiple comparison test was performed for data analysis. ***: P≤0.001 and ****: P≤0.0001.

### NLRP3 activating stimuli disrupt endosome trafficking

We next set out to test the hypothesis that NLRP3-activating stimuli disrupt endosome to TGN cycling that could explain the observed redistribution of TGN38/46. To assess the effects of NLRP3-activating stimuli and monensin on endosome to TGN trafficking we used fluorescently-labelled Shiga toxin B-subunit (STxB-Cy3) as described previously (*15*). STxB is internalised to early endosomes from where it undergoes retrograde transport to the TGN *en route* to the endoplasmic reticulum (Fig. 3A). HeLa cells were incubated with NLRP3-activating stimuli for 30 minutes at 37°C before incubation with STxB-Cy3. Trafficking to the TGN was assessed after 20 and 45 minutes by measuring STxB-Cy3 co-localization with Golgin-97 (Fig. 3B&C, fig S4A). All stimuli appeared to reduce co-localization of STxB-Cy3 and Golgin-97 at 20 minutes, but this was only found to be quantitatively significant following treatment with nigericin or imiquimod (Fig. 3B). At 45 minutes, there was increased co-localization with STxB-Cy3 and Golgin-97 in vehicle-treated cells, which was significantly reduced following monensin or nigericin treatment (Fig. 3B&C). This indicates that endosome to TGN trafficking is impaired by NLRP3 inflammasome-activating stimuli (Fig. 3B&C).

**Fig. 3.**
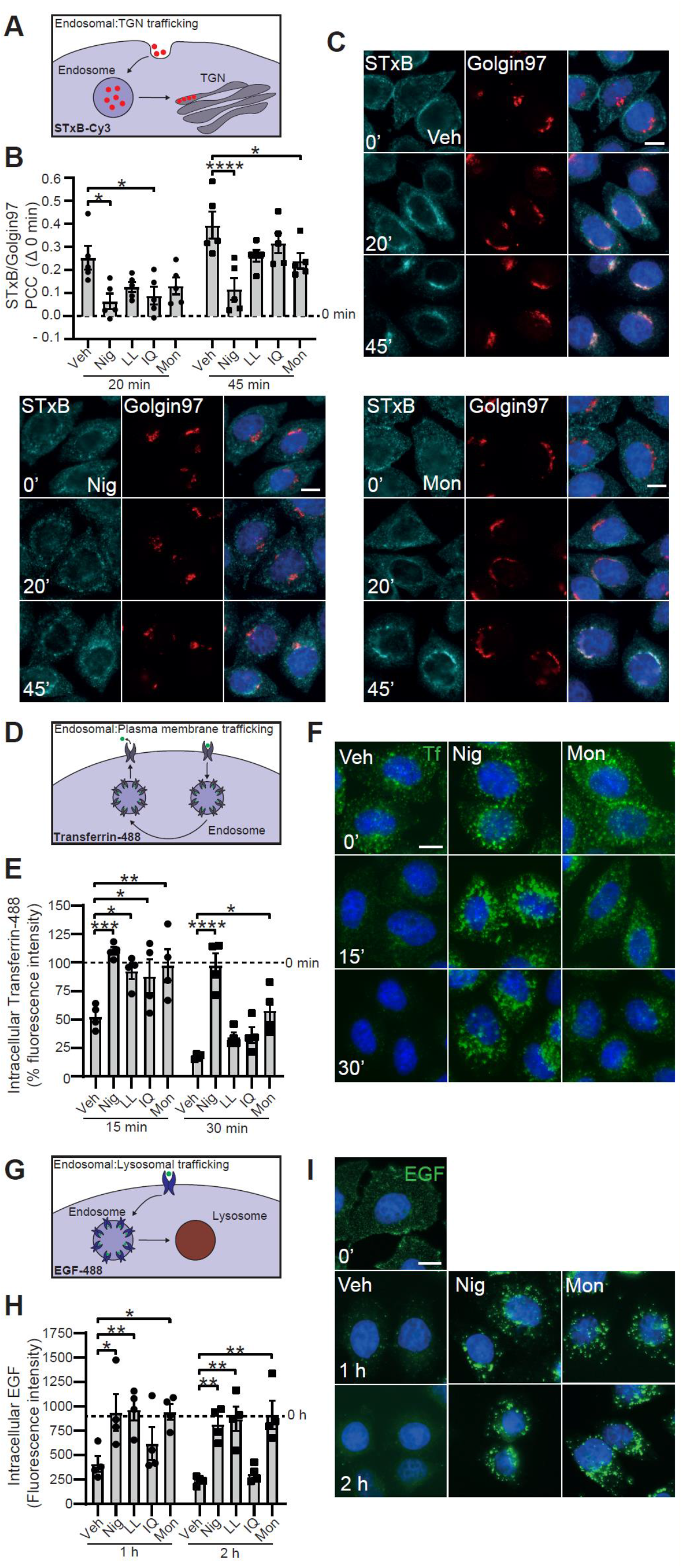
Endosomal trafficking is disrupted by NLRP3-activating stimuli. **(A)** Schematic of shiga toxin B (STxB) trafficking. STxB is endocytosed and trafficked from the plasma membrane to the TGN via endosomes. **(B)** HeLa cells were pretreated with vehicle (Veh), nigericin (Nig 10 μM), LLOMe (LL, 1 mM), imiquimod (IQ, 75 μM) or monensin (Mon, 10 μM) prior to loading of STxB-Cy3 (0.4 μg mL^−1^). Quantification of STxB-Cy3 trafficking to the TGN was measured by the change in Pearson’s correlation coefficient (PCC) between STxB-Cy3 and Golgin-97 at 20 and 45 min (compared to 0 min) following STxB-Cy3 addition (n = 5). **(C)** Representative immunofluorescence images of STxB-Cy3 (cyan) and Golgin-97 (red) in HeLa cells treated with vehicle, nigericin or monensin from (B). **(D)** Schematic of Transferrin trafficking. Transferrin binds to the transferrin receptor and is endocytosed into early endosomes, where it releases bound iron, before cycling back to the plasma membrane to be released extracellularly. **(E)** HeLa cells were treated with transferrin-488 (5 μg mL^−1^) and with either vehicle, nigericin, LLOMe, imiquimod, or monensin for 30 min before replacement with media containing unlabelled transferrin and the respective stimulus. Quantification of remaining cell-associated transferrin-488 was measured by the fluorescence intensity of transferrin-488 at 15 and 30 min following removal of extracellular transferrin-488 (n = 4). **(F)** Representative immunofluorescence images of transferrin-488 (Tf, green) in HeLa cells treated with vehicle, nigericin or monensin from (E). **(G)** Schematic of EGF trafficking. EGF binds to the EGF receptor, which is endocytosed and trafficked through the endo-lysosomal pathway resulting in degradation of EGF. **(H)** Hela cells were incubated with EGF-488 (0.4 μg mL^−1^) on ice for 1 h (to allow surface binding of EGF-488) prior to removal of extracellular EGF-488 and treatment with vehicle, nigericin, LLOMe, imiquimod or monensin at 37°C. Quantification of EGF-488 degradation was measured by the fluorescence intensity of EGF-488 at 1 and 2 h following incubation at 37°C (n = 4). **(I)** Representative immunofluorescence images of EGF-488 (green) in HeLa cells treated with vehicle, nigericin or monensin from (G). Scale bars: 10 μm. Values are mean ± SEM. Unmatched two-way ANOVA followed by Sidak’s multiple comparison test (compared to vehicle) were performed for data analysis. *: P≤0.05, **: P≤0.01, ***: P≤0.001, ****: P≤0.0001.

We then went on to investigate whether NLRP3-activating stimuli and monensin caused a more general disruption of endosome trafficking by looking at other endosome trafficking steps. Iron-loaded transferrin is endocytosed *via* clathrin-coated vesicles into early endosomes, where it releases bound iron due to the acidic pH, before cycling back to the plasma membrane, either directly or *via* recycling endosomes, and release into the extracellular space (*16, 17*) (Fig. 3D). We used a modified version of a previously described transferrin recycling assay (*18*) to determine the effects of NLRP3-activating stimuli. HeLa cells were incubated with fluorescently-conjugated transferrin (Tf-488) in the presence of NLRP3-activating stimuli or monensin for 30 minutes to allow both transferrin uptake and NLRP3 activation. These cells then underwent an acid wash on ice to remove cell surface transferrin and were either fixed immediately, to assess levels of transferrin endocytosis, or incubated for a further 15 or 30 minutes in the presence of excess unlabelled apo-transferrin to assess recycling. As shown in Fig. 3E and F (and fig. S4B&C), none of the NLRP3-activating stimuli tested, nor monensin, affected transferrin uptake (fluorescence at time 0, fig. S4B). In contrast, all treatments caused a retention of transferrin within the cell at 15 minutes (Fig. 3E&F, fig. S4C). This effect on transferrin cycling was most apparent for nigericin and monensin, although all stimuli caused decreased cycling (Fig. 3E&F), indicating that all NLRP3 activating stimuli tested, and monensin, perturbed endosomal recycling to the plasma membrane.

We next investigated the effects of NLRP3 activating stimuli on trafficking from endosomes to lysosomes. To investigate endosome to lysosome trafficking we utilised an established Epidermal Growth Factor (EGF) trafficking assay (*18, 19*) (Fig 3G). HeLa cells were incubated with fluorescently-labelled EGF (EGF-488) on ice for 1 hour to allow binding of EGF to the EGF receptor at the plasma membrane. EGF uptake and trafficking were initiated by shifting the cells to 37°C, and EGF delivery to the degradative lysosomal compartment assessed by loss of cellular EGF-488 fluorescence. Intracellular EGF fluorescence was therefore assayed 1 and 2 hours post-stimulation with NLRP3-activating stimuli or monensin by fluorescence microscopy (Fig. 3H&I, fig. S4D). All treatments except imiquimod significantly prevented the degradation of EGF, suggesting that they had prevented trafficking to the lysosomes (Fig. 3H&I).

These data show that the NLRP3-activating stimuli nigericin, LLOMe, and imiquimod perturb endosomal trafficking pathways with imiquimod having the mildest effect of stimuli tested. Thus, all NLRP3-activating stimuli investigated perturbed endosome trafficking to some extent.

### Effects of perturbing endosomal trafficking on NLRP3 activation

The data so far indicate a disruption of endosomal trafficking by NLRP3-activating stimuli. The fact that monensin can disrupt endosome cycling and not act as a potent NLRP3 activating stimulus suggests that while perturbed endosomal trafficking may be necessary for NLRP3 inflammasome activation, it is not sufficient. To explore this further we hypothesised that the effects of NLRP3 activating stimuli could be potentiated by disrupting endosome trafficking. To this point in our experiments, imiquimod had caused the mildest perturbation of endosomal trafficking of the NLRP3-activating stimuli tested. We thus considered the possibility that we could potentiate the effects of imiquimod by firstly perturbing endosome trafficking using monensin. To do this, LPS-primed mouse primary BMDMs were treated with monensin for 2 hours before adding imiquimod for 2 hours. We then assayed IL-1β release and cell death. Monensin pre-treatment of LPS-primed BMDMs greatly potentiated imiquimod-induced IL-1β release and cell death (Fig. 4A&B), ASC oligomerisation (measured as described previously (*20*)) (Fig. 4C), and processing of caspase-1 and gasdermin D (Fig, 4D). The exacerbation of IL-1β release and cell death caused by monensin pre-treatment were NLRP3-dependent as they were blocked by the specific NLRP3 inhibitor MCC950 (Fig. 4E&F). To study the effects of monensin pre-treatment on ASC speck formation further, we used immortalized mouse BMDMs (iBMDMs) and iBMDMs stably expressing ASC labelled with the fluorescent protein mCherry (*21*). In WT iBMDMs, monensin enhanced imiquimod-dependent IL-1β release and cell death (Fig. 4G&H), and in ASC-mCherry iBMDMs, significantly enhanced NLRP3-dependent ASC oligomerisation and speck formation (Fig. 4I-K). In the absence of imiquimod, monensin had no effect on IL-1β release or ASC speck formation in LPS-treated cells. We examined the effects of these treatments on Golgin-97 and TGN38 in LPS-primed iBMDMs. In vehicle-treated cells both Golgin-97 and TGN38 were compact in the peri-nuclear space (fig. S5). Monensin treatment disrupted the localization of both Golgin-97 and TGN38 and caused a separation in their co-localization as seen above, consistent with effects on endosome to TGN cycling, and also fragmentation of the TGN (fig. S5). Imiquimod had minimal effects on Golgin-97 and TGN38 distribution, while in imiquimod treated cells that had undergone prior treatment with monensin the disrupted Golgin-97 and TGN38 distribution was maintained (fig. S5).

**Fig. 4.**
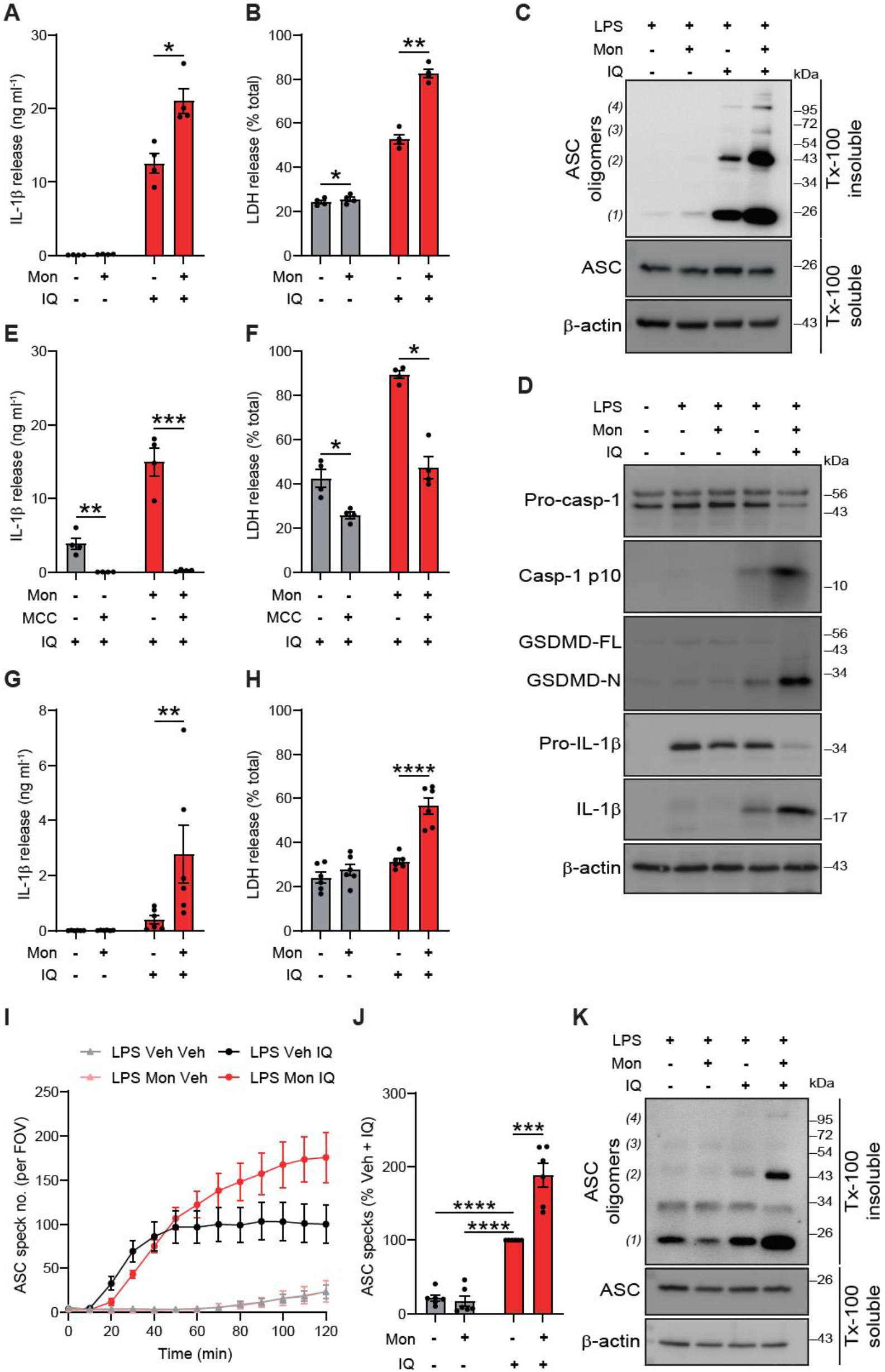
Monensin potentiates imiquimod-induced NLRP3 activation in primary and immortalized BMDMs. **(A-D)** LPS-primed primary BMDMs were treated with vehicle or monensin (Mon, 10 μM) for 2 h prior to imiquimod stimulation (IQ, 75 μM) for a further 2 h. Supernatants were assessed for (A) IL-1β and (B) LDH release (n = 4). Cell lysates and supernatants were assessed for (C) DSS-crosslinked ASC oligomers (n = 3), or (D) inflammatory protein content by western blotting (n = 4). **(E-F)** LPS-primed primary BMDMs were treated with vehicle or monensin (10 μM) for 2 h prior to MCC950 treatment (10 μM, 15 min) before subsequent imiquimod stimulation (75 μM) for a further 2 h. Supernatants were assessed for (E) IL-1β and (F) LDH release (n = 4). **(G-H)** LPS-primed WT iBMDMs were treated with vehicle or monensin (10 μM) for 1 h prior to imiquimod stimulation (75 μM) for a further 2 h. Supernatants were assessed for (G) IL-1β and (H) LDH release (n = 6). **(I-J)** LPS-primed iBMDMs stably expressing ASC-mCherry were treated with vehicle or monensin (10 μM) for 2 h prior to Ac-YVAD-CMK (100 μM, 15 min) followed by imiquimod stimulation (75 μM) for a further 2 h. (I) ASC speck formation was measured, and (J) ASC speck number at the final time point of 2 h is shown (% of veh + imiquimod treatment) (n = 6). **(K)** LPS-primed iBMDMs were treated with vehicle or monensin (10 μM) for 1 h prior to imiquimod stimulation (75 μM) for a further 2 h. Cell lysates and supernatants were assessed for DSS-crosslinked ASC oligomers (n = 4). IL-1β release was measured by ELISA. Values are mean ± SEM. Repeated-measures (A, B, E, F) or unmatched (G, H) two-way ANOVA followed by Sidak’s multiple comparison test, or one-sample t test versus a value of 100% followed by Holm-Sidak correction (J), were performed for data analysis. *: P≤0.05, **: P≤0.01, ***: P≤0.001, ****: P≤0.0001.

We also investigated whether the effects of nigericin were potentiated by prior treatment of LPS-primed iBMDMs with monensin. In this case monensin had no effect on nigericin-induced IL-1β release, or ASC speck formation in ASC-mCherry expressing iBMDMs (fig. S6), potentially as the effects of nigericin on endosomal trafficking and NLRP3 activation are already maximal.

### NLRP3 can localize to endosomal membranes

Considering our results suggesting a link between dysregulation of endosomal trafficking and NLRP3 stimulation, we next wanted to assess whether NLRP3 could localize to endosomal membranes under stimulatory conditions. For this purpose, we assessed NLRP3 localization in COS7 cells stably expressing low levels of NLRP3-mVenus. In the absence of stimulation, NLRP3-mVenus was localized predominantly in the cytosol and occasionally at the peri-nuclear TGN (Fig. 5A-C, fig. S7&S8). Stimulation with our panel of NLRP3-activating stimuli or monensin caused a redistribution of some of the NLRP3-mVenus to cytoplasmic puncta that partially co-localized with EEA1- and CD63-containing endosomes as well as LAMP1-containing lysosomes (Fig. 5A-C, fig. S8A-C). In contrast, NLRP3-mVenus puncta rarely co-localized with the trans-Golgi marker Golgin-97 (fig. S7). These data suggest that NLRP3 can localize to multiple endosomal membranes upon stimulation. Indeed a co-localization of TGN38, EEA1, and NLRP3-GFP in HeLa cells treated with nigericin was also observed previously (*5*).

**Fig. 5.**
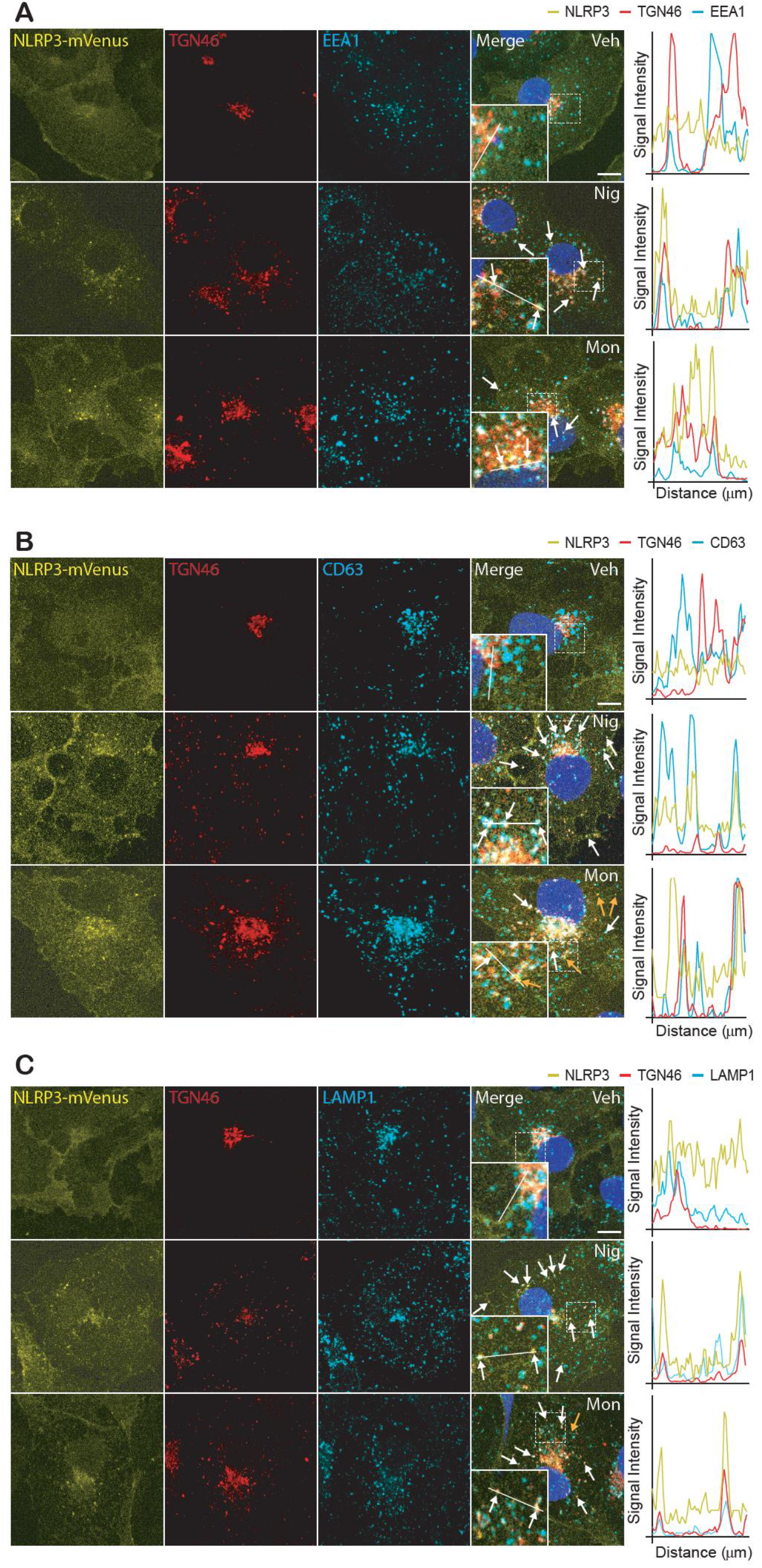
NLRP3-mVenus co-localizes with TGN46-, EEA1- and CD63-positive puncta after stimulation with NLRP3-activating stimuli. **(A-C)** Immunofluorescence images of COS7 cells stably expressing NLRP3-mVenus. Cells were stimulated for 90 min with vehicle (Veh, 0.5% ethanol (v/v)), nigericin (Nig, 10 μM) or monensin (Mon, 10 μM). Also shown are line graphs depicting changes in fluorescence intensity (min = 0, max = 250) over 10 μm, for each of the conditions and stains. Images are representative of 3-5 independent experiments. Blue represents nuclei staining by DAPI. White arrows indicate co-localization between NLRP3 and **(A)** EEA1-, **(B)** CD63- or **(C)** LAMP1-positive puncta; yellow arrows indicate NLRP3 puncta alone. Dashed lines highlight the areas depicted in the zoomed insets.

NLRP3 was previously reported to associate with the inositol lipid PtdIns4P (*5*). Using the PtdIns4P probe SidM-mCherry (*22*) transiently transfected into NLRP3-mVenus-expressing COS7 cells, we sought to determine any co-localization between NLRP3 and PtdIns4P. Under resting conditions NLRP3 was diffuse in the cytosol, whereas the PtdIns4P probe concentrated on the TGN, as expected (*22*) (Fig. 6A-D, fig. S9D&S10A). Upon treatment with nigericin, PtdIns4P was lost from the TGN and accumulated on discrete cytoplasmic puncta, many of which co-localized with NLRP3 (Fig. 6A-D). A similar effect was seen with LLOMe (fig. S9A,D, fig. S10A), whereas monensin did not affect distribution of the PtdIns4P probe, but did cause NLRP3 to localise to puncta as expected (Fig. 6A-D). In response to NLRP3-activating stimuli, the NLRP3-mVenus- and PtdIns4P-containing puncta showed a partial co-localization with markers of late endosomes (Fig. 6B&D, fig. S9B&D) and lysosomes (Fig. 6C&D, fig. S9C&D). Whilst monensin disrupted TGN46 trafficking, its failure to affect PtdIns4P localization could be part of an explanation for why it is not a sufficient stimulus for NLRP3 activation. As discussed previously, pre-treatment with monensin potentiated imiquimod-induced NLRP3 activation. Interestingly, imiquimod alone did not affect PtdIns4P distribution, whereas treatment with monensin and subsequently imiquimod caused accumulation of PtdIns4P on late endosomal and lysosomal compartments also containing NLRP3, albeit to a lesser extent than nigericin (fig. S9A-D). Interestingly, co-localization of PtdIns4P with NLRP3 and EEA1 after stimulation with NLRP3 activators was limited and only observed with nigericin (fig. S10).

**Fig. 6.**
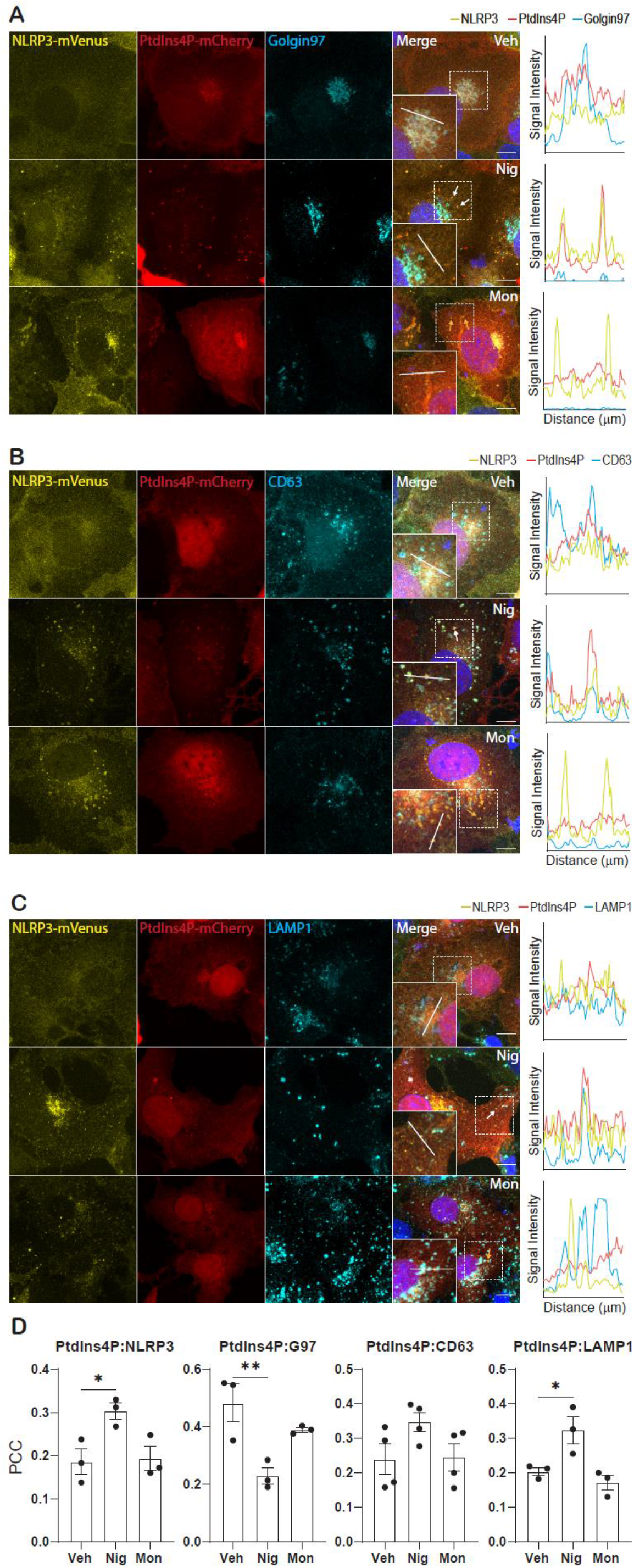
NLRP3 mVenus co-localizes with PtdIns4P on CD63- and LAMP1-positive puncta after stimulation with NLRP3-activating stimuli. **(A-C)** Fluorescence and immunofluorescence images of COS7 cells stably expressing NLRP3-mVenus. Cells were stimulated for 90 min with vehicle (Veh, 0.5% ethanol (v/v)), nigericin (Nig, 10 μM) or monensin (Mon, 10 μM). Co-localization between NLRP3-mVenus, PtdIns4P and **(A)** Golgin-97, **(B)** CD63, or **(C)** LAMP1 was then assessed. Also shown are line graphs depicting changes in fluorescence intensity (min = 0, max = 250) over 10 μm, for each of the conditions and stains. Images are representative of 3-4 independent experiments. Blue represents nuclei staining by DAPI. White arrow indicates co-localization between NLRP3, PtdIns4P, and **(A)** Golgin-97 **(B)** CD63- or **(C)** LAMP1 positive puncta; yellow arrows indicate NLRP3 puncta alone. Dashed lines highlight the areas depicted in the zoomed insets. **(D)** Pearson’s correlation coefficient (PCC) between PtdIns4P and the proteins indicated. Values are mean ± SEM, (n=3-4). One-way ANOVA followed by Dunnett’s multiple comparison test (versus vehicle) was performed for data analysis. *: P≤ 0.05, **: P≤0.01.

## Discussion

There is a broad literature linking organelle stress to the activation of the NLRP3 inflammasome. Following analysis and review of this literature we hypothesized that a cellular stress arising from disrupted endosomal cycling could serve as a trigger for NLRP3-inflammasome activation (*1*). Here, we established that NLRP3-activating stimuli cause TGN38/46 to become trapped in endosomes, which may account for at least some of the dispersed TGN38/46 observed in previous literature (*5*). Further, using established assays of endosome trafficking to the TGN, to the lysosomes, or to the plasma membrane (*18*), we established that all NLRP3-activating stimuli tested, to some extent, disrupted an aspect of endosome trafficking. These results suggest that a common effect of NLRP3-activating stimuli is a disruption of endosomal trafficking. Disrupted endosomal trafficking of TGN38/46 itself, however, appears insufficient to activate the NLRP3 inflammasome. Monensin, a well-established and potent disruptor of endosome trafficking (*7*), by itself had minimal effects on NLRP3 inflammasome activation, even at a concentration that profoundly affected endosomal trafficking of TGN38/46. These data suggest that if disruption of endosomal trafficking is important, it must occur concomitantly with something else to activate NLRP3. A possible candidate is the accumulation of PtdIns4P on endosomal and lysosomal membranes, given that NLRP3-activating stimuli cause this effect whereas monensin does not.

The NLRP3 inflammasome activating stimulus imiquimod was included in our panel as, in contrast to the majority of canonical NLRP3 activating stimuli, its activation of NLRP3 occurs in the absence of K^+^ ion efflux (*10*). In all assays on endosomal trafficking investigated, imiquimod had the mildest effect, though its trapping of TGN46 in endosomes was significant. When we fully disrupted endosomal trafficking by treating LPS-primed macrophages with monensin, and then treated with imiquimod we potentiated NLRP3-inflammasome activation and IL-1β release. These data highlight how manipulation of a potential endosome stress-sensing pathway interacts with an aspect of NLRP3 biology to maximise an inflammatory response. That monensin did not potentiate nigericin-induced IL-1β release may reflect that nigericin alone is already a very efficient disruptor of endosome trafficking.

Previous research has shown a co-localization of NLRP3 inflammasomes with TGN38/46 positive structures (*5*). Research in endothelial cells shows NLRP3 co-localizes to endosomes with complement membrane attack complexes (*23*). In macrophages, caspase-1 cleaves EEA1, and active caspase-1 co-localizes to EEA1-positive endosomes in cells treated with hypotonicity, a NLRP3-activating stimulus (*24*). Furthermore, NLRP3 can interact with PtdIns4P via a conserved polybasic motif (*5*), and endosomal membranes are known to contain this lipid (*22*). In our experiments, PtdIns4P was lost from the TGN and accumulated on CD63-positive late endosomes and LAMP1-positive lysosomes in response to NLRP3-activating stimuli. PtdIns4P on CD63- and LAMP1-positive compartments also co-localized with NLRP3. However, monensin alone did not cause a change in PtdIns4P distribution, despite inducing the formation of NLRP3 puncta. Imiquimod had the mildest effect upon the PtdIns4P, but when combined with monensin, PtdIns4P and NLRP3 were both redistributed similar to other NLRP3 agonists. It is possible that NLRP3 recruitment to, and concentration at, endolysosomal membranes is important for its activation, however it cannot be sufficient considering that monensin alone fails to activate NLRP3. Our results suggest PtdIns4P accumulation on endolysosomal membranes is important in the activation process, but additional events such as post-translational modification of NLRP3, which could also occur on the membrane, may also be necessary. Many signalling processes are known to occur on endosomal membranes (*11*), and inflammasome signalling may represent yet another. Whether NLRP3 persists on the membrane following its activation, or dissociates, is the subject of future research. Thus our data, and other observations from the literature, support the possibility that NLRP3 senses a disruption of endosome trafficking and that its enrichment on membranes of endosomal origin is important for activation of the NLRP3 inflammasome.

Understanding the activation of the NLRP3 inflammasome remains an enduring question in the field. Our data consolidate observations in the literature that disruption of endosomal trafficking could contribute to the stress sensed by NLRP3 that is, in part, required for its activation, and that the endosomes are viable candidates for a sub-cellular site of importance to inflammasome activation. Understanding these mechanisms further will reveal new insights into NLRP3 biology that may identify new ways of targeting NLRP3-dependent inflammation in disease.

## Author contributions

Author contributions are as follows:

Conceptualization: DW, ML, DB, GLC
Methodology: AA, HB, GLC,
Investigation: BL, DW, FMS, RW, CH, PS, JG
Visualization: BL, FMS, RW, CH, PS, JG, DB, ML, GLC
Funding acquisition: DB, ML, GLC
Project administration: DB, ML, GLC
Supervision: DB, ML, GLC
Writing – original draft: DB, ML, GLC
Writing – review & editing: DB, ML, GLC, BL, JG, CH, PS, RW,

We are also grateful to Professor Philip Woodman of the University of Manchester for useful advice and guidance on endosome trafficking assays and to the Bioimaging and Flow Cytometry Core Facilities.

## Data and materials availability

All data will be made available upon reasonable request to the principal investigator and bespoke or unique materials used will be made available via a MTA with the University of Manchester.

