## Supplementary files for "NLRP3 activation in response to disrupted endocytic traffic"

### Supplementary Materials

### Materials and Methods

#### *Reagents*

Transferrin from Human Serum, Alexa Fluor™ 488 Conjugate (T13342) and epidermal growth factor (EGF) Alexa Fluor™ 488 (E13345) were from Thermo-Fisher. Human holo-transferrin (T4132) was from Sigma-Aldrich. Shiga Toxin B-Cy3 was kindly provided by Prof. Ludger Johannes (Curie Institute, Paris).

Lipopolysaccharide (LPS, *Escherichia coli 026:B*6), nigericin (N7143) and MCC950 (PZ0280) were purchased from Sigma. L-Leucyl-L-Leucine methyl ester (LLOMe, 16008-1) was from Cayman Chemical, imiquimod (R837) from InvivoGen, and monensin sodium salt (5223/500) from Tocris Bioscience. Ac-YVAD-CMK (4018838) was from Cambridge Bioscience.

Specific antibodies were used for immunocytochemistry targeting: TGN38 (0.1 μg $mL^{-1}$, AHP499G, Bio-Rad), TGN46 (0.25 μg $mL^{-1}$, AHP500GT, Bio-Rad), Golgin-97 (2 μg $mL^{-1}$, ab84340, abcam), EEA1 (2.5 μg $mL^{-1}$, 610457, BD biosciences; 1:100 dilution of stock, MA5-14794, ThermoFisher), CD63 (1 μg $mL^{-1}$, CBL553, Sigma-Aldrich), LAMP1 (1:300 dilution of stock, MA1-164, Thermo-Fisher, or 9091, CST) and MICAL-L1 (1:100 dilution of stock, H00085377-B01P, abnova). Alexa Fluor secondary antibodies (488, 594, and 647) were from Thermo-Fisher. For western blotting, goat anti-mouse IL-1β (AF-401-NA) was from R&D Systems, rabbit anti-mouse ASC (67824) was from CST, rabbit anti-mouse caspase-1 (ab179515) and rabbit anti-mouse gasdermin D (ab209845) were from Abcam, and rabbit anti-goat IgG (P044901-2) and goat anti-rabbit IgG (P044801-2) were from Agilent. Anti-β-actin-peroxidase (A3854) was from Sigma.

#### *Cell Culture*

HeLa and COS-7 (ATCC) cells were cultured in T75 flasks in DMEM supplemented with 10% FBS, 100 U $mL^{-1}$ penicillin, 100 μg $mL^{-1}$ streptomycin, and 1 mM sodium pyruvate. Cells were seeded on coverslips in a 24 well plate (Corning) at $1x10^5$ $ml^{-1}$ density and left to adhere overnight at 37 °C and 5% $CO_2$.

iBMDMs were originally a gift from Professor Clare Bryant (Department of Veterinary Medicine, University of Cambridge) and were modified to express ASC-mCherry in Manchester *(21)*. WT and ASC-mCherry iBMDMs were cultured in DMEM supplemented with 10% FBS,

100 U $mL^{-1}$ penicillin, 100 μg $mL^{-1}$ streptomycin and 1 mM sodium pyruvate, seeded at a density of 7.5 x $10^5$ $mL^{-1}$, and left to adhere overnight at 37 °C and 5% $CO_2$.

Primary mouse BMDMs were prepared from the femurs of 3–6 month-old male and female C57BL/6 mice. Bone marrow was harvested, red blood cells were lysed and the resulting cells were passed through a cell strainer and cultured in 70% DMEM (containing 10% FBS, 100 U $mL^{-1}$ penicillin, 100 μg $mL^{-1}$ streptomycin, and 1 mM sodium pyruvate), supplemented with 30% L929 mouse fibroblast-conditioned medium for 7 days at 37 °C and 5% $CO_2$. Cells were seeded at a density of 1 x $10^6$ $mL^{-1}$ and left to adhere overnight.

Lenti-X 293T cells (#632180, Takara) were cultured in DMEM supplemented with 10% FBS, 100 U $mL^{-1}$ penicillin, 100 μg $mL^{-1}$ streptomycin and 1 mM sodium pyruvate.

***Plasmids***

A 3rd generation lentiviral EF1a-NLRP3-mVenus expression vector was generated by HiFi assembly (NEB). Briefly, a custom built Lenti-EF1a-Multiple cloning site-pGK-Puro vector was digested with KpnI and NotI (NEB) and column purified (Bioline). Hifi assembly fragments were amplified, using HF KOD polymerase, from NLRP3 (fragments 1 and 2) or mVenus (fragment 3) template DNA using the primers Frag 1f actagtccagtgtggtggtaccggtggatccgccaccatgaagatggcaagcacccg and Frag 1r tcgtcaaaggcaccttgcagctcatcgaagccgtccatgag, Frag 2f tcatggacggcttcgatgagctgcaaggtgcctttgacg, and Frag 2r gcagcggagccagcggagccccaagaaggctcaaagacgacg, Frag 3f tcgtctttgagccttcttggggctccgctggctcc and Frag 3r aacagatggctggcaactagaaggcacaggcggccgctcacttgtacagctcgtccatgc, assembled into the backbone vector according to manufacturer's instructions, transformed into NEB C3040 cells. Clones were sequenced verified before being taken forward for lentiviral packaging. Vectors generated in this study will be submitted to Addgene. Packaging plasmids pMD2.G and psPAX2 for lentivirus production were a gift from Didier Trono (Addgene plasmid#12259 and #12260)

***Virus production and generation of COS-7 cells stably expressing NLRP3-mVenus.***

Lenti-X 293 cells were plated (1 x $10^6$ per well in a six-well plate) 24h before the transfection with 1.2 μg pMD2.G, 0.4 μg psPAX2 and 1.5 μg of NLRP3-Venus using Lipofectamine 2000 (Invitrogen). Transfection was performed following the Lipofectamine manufacturer's instructions. 24h after transfection the media was replaced by fresh media and cells were further incubated for 2 days. Supernatants were then collected and filtered with a 0.45μm filter to obtain a cell free extract of viral particles, which was used to transduce COS-7 cells.

For the transduction, 5x$10^4$ COS-7 cells plated the day before, were incubated for 7h with a mix of 8 μg $mL^{-1}$ polybrene (Sigma) and the viral particles. After 48h virus containing media was removed and positive expressing cells were selected with puromycin 1.5 μg $mL^{-1}$ for 10 days. After this cells were FAC sorted to select a cell population homogeneously expressing NLRP3-Venus.

***Transient Transfection***

The day before transfection, COS-7 stably expressing NLRP3-mVenus cells were seeded at $7x10^4$ cells $ml^{-1}$ in a 24-well flat-bottomed plate (Corning) and left to adhere overnight at 37°C with 5% $CO_2$. Cells were transfected using the Lipofectamine 3000 kit (Invitrogen) according to manufacturer's instructions. Cells were transfected with 300ng SidM-mCherry. Whilst the DNA-lipid complex was incubating, cell media was replaced with Opti-MEM (Thermo-Fisher). DNA-lipid complex was then added to cells and left to incubate for 20-22 hours at 37°C, 5% $CO_2$.

#### *Cell stimulation*

For stimulation of COS-7 and HeLa cells, culture media was replaced with fresh serum-free DMEM before stimulation with vehicle (ethanol 0.5% v/v), nigericin (10 µM), LLOMe (1 mM), imiquimod (75 µM) or monensin (10 µM) for 90 min.

For stimulation of primary BMDMs, cells were first primed with either vehicle or LPS (1 µg $mL^{-1}$) for 4 h. The culture media was then replaced with fresh serum-free media with and without MCC950 (10 µM). After 15 min cells were stimulated with vehicle (ethanol 0.5% v/v), nigericin (10 µM), LLOMe (1 mM), imiquimod (75 µM) or monensin (10 µM) for 90 min.

*NLRP3 potentiation experiments*

Macrophages were primed with LPS (1 µg $mL^{-1}$) for 4 h (primary BMDM) or 2 h (iBMDM) prior to treatment with either vehicle (ethanol 0.5% v/v) or monensin (10 µM), for 1 or 2 h in serum-free DMEM. In some experiments, MCC950 (10 µM) was then added for 15 min. Imiquimod (75 µM, 2 h) or nigericin (10 µM, 1 h) was then added to activate NLRP3. At the end of the experiment, supernatants were collected for further analysis, or cells were prepared for western blotting or immunocytochemistry.

#### *Endosome trafficking assays*

*Shiga toxin trafficking*

HeLa cells were seeded out at a density of $0.25x10^6$ $mL^{-1}$ on 13 mm glass coverslips the day before experiments. HeLa cells were incubated in serum free DMEM with NLRP3 activating stimuli (nigericin 10 µM, LLOMe 1 mM, imiquimod 75 µM), monensin (10 µM), or vehicle (ethanol, 0.5% v/v) for 30 min at 37°C. Cells were washed once with ice cold PBS before incubation with shiga toxin B-cy3 (0.4 µg $mL^{-1}$ in 2 mg $mL^{-1}$ BSA in PBS) on ice for 30 min. Cells were washed twice in ice cold PBS and then either fixed with PFA (4% w/v) (for a 0 min time point) or incubated in serum free DMEM with NLRP3 activating stimuli for a further 20 or 45 min at 37°C before PFA fixation. Fixed HeLa cells were incubated overnight at 4°C with specific antibodies targeting Golgin-97.

*Transferrin trafficking*

HeLa cells were washed three times in serum free DMEM and incubated for 30 min in DMEM containing Transferrin Alexa Fluor-488 (5 µg $mL^{-1}$) and NLRP3 activating stimuli (nigericin 10 µM, LLOMe 1 mM, imiquimod 75 µM), monensin (10 µM) or vehicle (ethanol, 0.5% v/v). Cells underwent a 3 min ice cold acid wash (50 mM glycine, 0.1 M NaCl in dH20, pH=3) before being washed three times in serum free media containing holo-transferrin (0.1 mg $mL^{-1}$). Cells were either fixed at this point (0 min) or incubated in serum free DMEM containing unlabelled holo-transferrin (0.1 mg $mL^{-1}$) for 15 or 30 min at 37 ºC before fixation. Cells were fixed on coverslips in 4% paraformaldehyde (PFA).

*EGF trafficking*

HeLa cells were washed three times with PBS before incubation with EGF Alexa Fluor-488 (0.4 µg $mL^{-1}$) in serum free DMEM (supplemented with 2% w/v BSA) on ice for 1 h. Cells were washed twice with cold PBS to remove unbound EGF-488. Cells were then either fixed with 4% PFA (for a 0 min time point), or stimulated with either a vehicle (ethanol 0.5% v/v) nigericin (10 µM), LLOMe (1 mM), imiquimod (75 µM), or monensin (10 µM) in serum free DMEM at 37ºC for 1 or 2 h before fixation (4% PFA).

***Immunocytochemistry***

Cells were fixed with 4% paraformaldehyde for 20 min before being permeabilized with PBST (0.1% Triton X-100) for 5 min at RT. To reduce nonspecific binding of antibodies the coverslips were incubated with blocking solution (5% BSA in PBST) for 30 min at RT, before incubation with primary antibodies diluted in blocking solution followed by Alexa Fluor secondary antibodies. Nuclei were stained using DAPI. The coverslips were then mounted on top of microslides using ProLong Gold antifade mounting reagent (Invitrogen) and left to dry overnight at RT.

***Fluorescence microscopy and image analysis***

*Widefield*

Widefield fluorescence microscopy was used for endosomal trafficking assays using STxB-Cy3, transferrin-488 and EGF-488. Images were captured on a Zeiss Axioimager D2 upright microscope using a 63× / 1.4 Plan Apochromat objective with a Coolsnap HQ2 camera (Photometrics) and Micromanager software (v1.4.23). Specific band pass filter sets were used to prevent bleed through from one channel to the next. Images were taken of 3-5 independent fields of view per treatment per time point within each replicate.

*Confocal*

1024 x 1024 images were captured using a 63× / 1.40 HCS PL Apo objective on a Leica TCS SP8 AOBS inverted or upright confocal microscope with LAS X software (v3.5.1.18803). The blue diode with 405 nm and the white light laser with 488 nm and 594 nm laser lines were used, with hybrid and photon-multiplying tube detectors with detection mirror settings of 415-478 nm (DAPI), 498-584 nm (FITC) and 604-750 nm (Texas Red), respectively. To prevent crosstalk between channels, images were acquired sequentially. Z-stacks were acquired with 0.3 µm steps between Z sections. Camera gain and exposure times were unchanged within each replicate. Images were taken of 3-5 independent fields of view per treatment per time point within each replicate with an average of 100 cells imaged for each condition.

*Image analysis*

Following image collection, image analysis was carried out using Fiji (Image J) software. The Pearson’s correlation co-efficient (PCC) for each condition was calculated using the Coloc 2 plugin. Maximum intensity projections were used for confocal images. For PCC analysis of

STxB, PCC at 20 and 45 minutes was normalised against the PCC at 0 minutes for each respective stimulus to determine change in PCC over time.

For fluorescence analysis of transferrin-488 and EGF-488 trafficking, the fluorescence intensity of 7-10 cells per field of view per experiment was measured. Background was manually calculated by taking independent measurements within each image from areas where cells were absent. The averaged background value was then subtracted from its respective mean fluorescence read-out. For transferrin-488, mean cell fluorescence was converted into percentage change against 0 minutes for each stimulus.

Fluorescence intensity line graphs were created using image J software. Fluorescence intensity of each channel was plotted individually over a 10 µm line then combined for comparison between channels. Line graphs depict fluorescence intensity between 0 and 250 grey value (a.u).

#### *ELISA*

Supernatants were assessed for IL-1β content by enzyme-linked immunosorbent assay (ELISA; DY401, DuoSet, R&D systems) according to manufacturer's instructions.

#### *LDH release assay*

Cell death was determined by measuring LDH release into the supernatant using a CytoTox 96 Non-Radioactive Cytotoxicity Assay (G1780, Promega) according to manufacturer's instructions.

#### *ASC speck formation live imaging*

iBMDMs stably expressing ASC-mCherry *(21)* were seeded overnight at a density of 7.5 x $10^5$ $mL^{-1}$ in black-walled 96-well plates. Cells were primed with LPS (1 µg $mL^{-1}$) for 2 h prior to treatment with either vehicle (ethanol 0.5% v/v) or monensin (10 µM), for 2 h in optimem. Ac-YVAD-CMK (100 µM) was added to all wells with or without MCC950 (10 µM) for 15 min, and imiquimod (75 µM) or nigericin (10 µM) was then added to activate NLRP3. Image acquisition began immediately after the addition of imiquimod or nigericin. Fluorescent images were acquired in duplicate every 10 min for 2 h using an IncuCyte S3 Live-Cell Analysis system (Essen Bioscience) at 37 °C using a 20X/0.61 S Plan Fluor objective and the red filter set for fluorescent images. Speck numbers were quantified manually using Fiji (ImageJ).

#### *ASC oligomerisation assay*

Primary BMDMs were seeded overnight at a density of 1 x $10^6$ $mL^{-1}$ in 12-well plates. Cells were primed with LPS (1 µg $mL^{-1}$) for 4 h prior to treatment with either vehicle (ethanol 0.5% v/v) or monensin (10 µM), for 2 h in serum-free DMEM. Imiquimod (75 µM, 2 h) was then added to activate NLRP3. Cells were then lysed in-well by direct addition of protease inhibitor cocktail and Triton-X-100 (1% v/v) into the culture medium. The combined cell lysate and supernatant was centrifuged at 6800 *x g* for 20 min at 4°C to generate Triton-X-100 soluble and insoluble fractions, followed by disuccinimidyl suberate (DSS)-crosslinking (2 mM, 30 min) of the insoluble fraction at RT, as described previously *(20)*. The insoluble fraction was subsequently centrifuged at 6800 x g for 20 min at 4°C, and then resuspended in 1X Laemmli buffer and heated at 95°C. The Triton-X-100 soluble fraction was concentrated by adding tricholoracetic acid (10% w/v), centrifuging at 18600 *x g* for 10 min at 4°C, washing the pellet with acetone, centrifuging again at 18600 *x g* for 10 min at 4°C before resuspending in 2X Laemmli buffer and heating at 95°C.

#### *Western blotting*

BMDMs were lysed in-well by direct addition of protease inhibitor cocktail and Triton-X-100 (1% v/v) into the culture medium at the end of the experiment. The combined cell lysate and supernatant was concentrated using tricholoracetic acid precipitation, as described above. Equal volumes of concentrated in-well lysates were loaded into 12% or 15% SDS-polyacrylamide gels. Samples were run and then transferred at 25 V onto nitrocellulose or PVDF membranes using a Trans-Blot® Turbo Transfer™ System (Bio-Rad). Membranes were blocked in milk (5% w/v) or BSA (2.5% w/v) in PBS Tween (0.1% v/v) for 1 h at RT. Membranes were incubated at 4°C overnight with goat anti-mouse IL-1β (0.25 μg $mL^{-1}$), rabbit anti-mouse ASC (0.101 μg $mL^{-1}$), rabbit anti-mouse caspase-1 (1.87 μg $mL^{-1}$) or rabbit anti-mouse gasdermin D (0.6 μg $mL^{-1}$) primary antibodies in 0.1% (IL-1β) or 2.5% (ASC, caspase-1, gasdermin D) BSA in PBS Tween). The membranes were washed with PBS Tween and incubated with rabbit anti-goat IgG (500 ng $mL^{-1}$, 5% milk in PBS Tween) or goat anti-rabbit IgG (250 ng $mL^{-1}$, 2.5% BSA in PBS Tween) at RT for 1 h. Proteins were visualised with Cytiva Amersham ECL Prime Western Blotting Detection Reagent (GE Healthcare, RPN2236) and G:BOX (Syngene) and Genesys software. β-Actin was used as a loading control.

#### *Data presentation and analysis*

Pearson's correlation coefficient, fluorescence intensity, ELISA, ASC speck, and cell death data are presented as mean ± standard error of the mean (SEM) with individual data points shown. Data were analyzed using unmatched or repeated measures one-way or two-way analysis of variance (ANOVA) with Sidak's or Dunnett's post-hoc test. Data normalized as a percentage were analyzed using a one-sample t test versus a value of 100%, followed by Holm-Sidak correction. All analyses were performed using GraphPad Prism (v8). Data were transformed prior to analysis where necessary to ensure equal variance between groups. Blots are representative of 4 experiments.

### Supplementary Figures

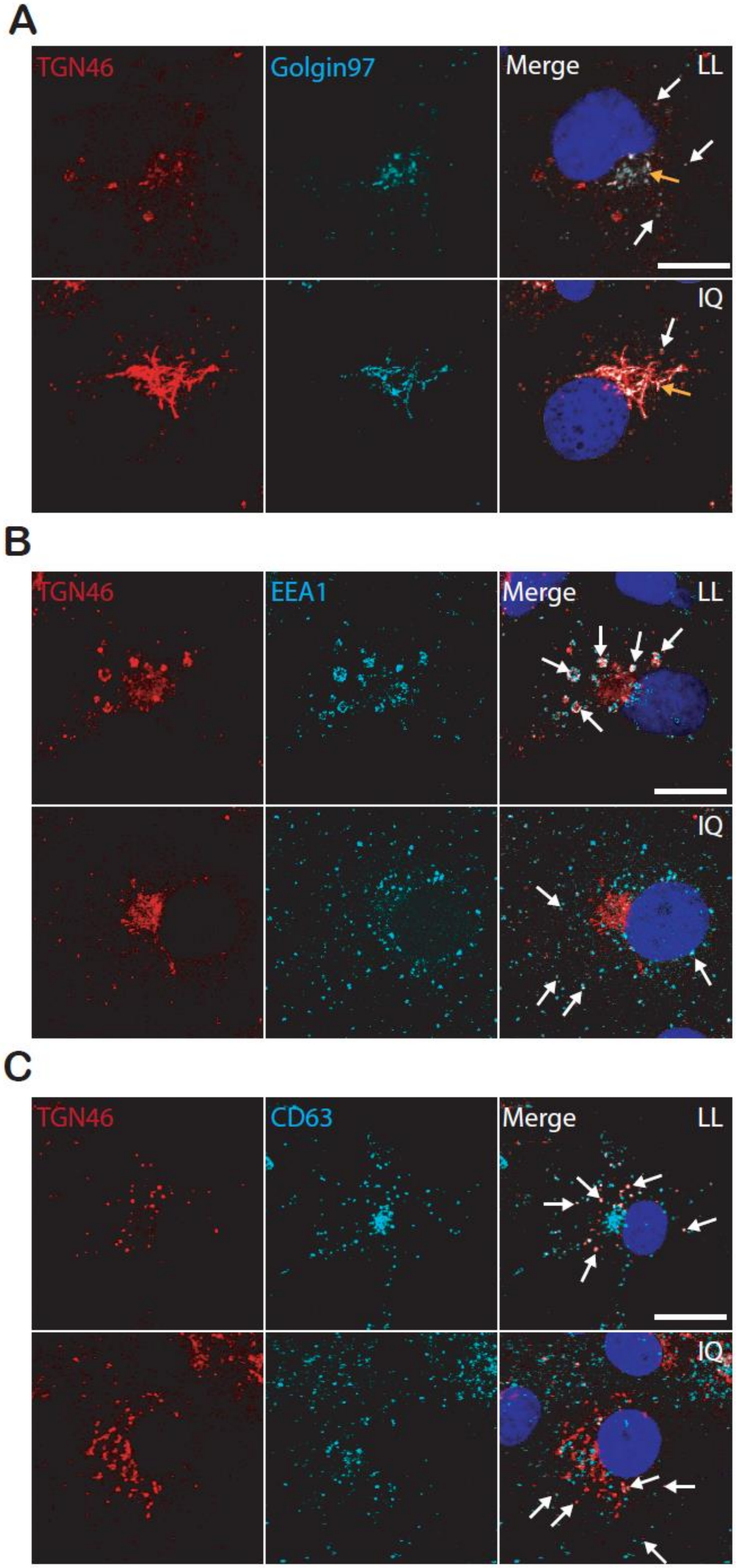

**Fig. S1. TGN46 co-localizes with endolysosomal membrane markers after stimulation of COS7 cells with NLRP3-activating stimuli.** COS7 cells were stimulated for 90 min with LLOMe (LL, 1 mM) or imiquimod (IQ, 75 µM). **(A-C)** Immunofluorescence images showing the co-localization of TGN46 with **(A)** Golgin-97, **(B)** EEA1, or **(C)** CD63. See also Fig 1. Images are representative of 3-5 independent experiments. Blue represents nuclei staining by DAPI. Yellow arrowheads indicate instances of co-localization at the Golgi, white arrowheads indicate co-localization on puncta. Scale bars: 10 µm.

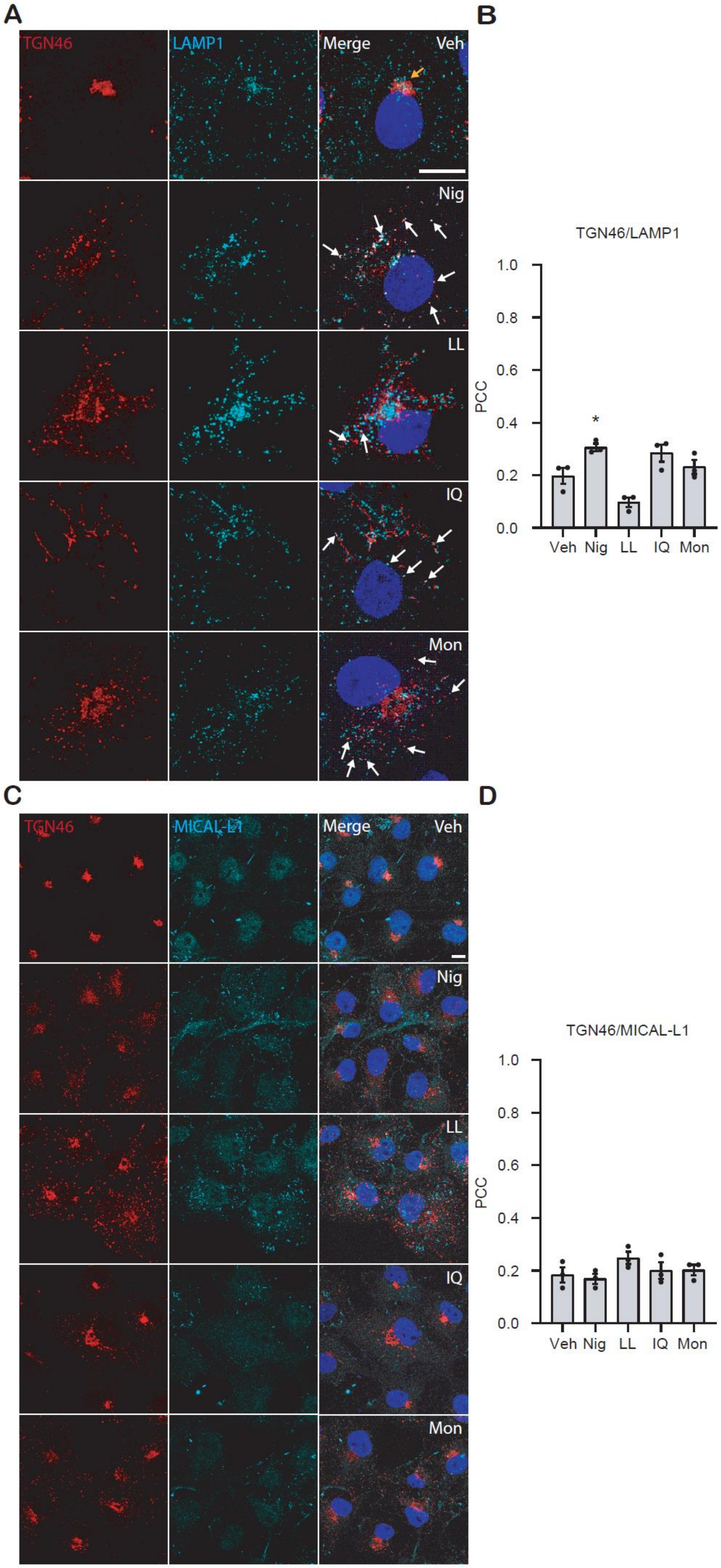
A
TGN46
LAMP1
Merge
Veh
Nig
LL
IQ
Mon
B
TGN46/LAMP1
PCC
1.0
0.8
0.6
0.4
0.2
0.0
*
Veh
Nig
LL
IQ
Mon
C
TGN46
MICAL-L1
Merge
Veh
Nig
LL
IQ
Mon
D
TGN46/MICAL-L1
PCC
1.0
0.8
0.6
0.4
0.2
0.0
Veh
Nig
LL
IQ
Mon

**Fig. S2. TGN46 can partially co-localize with lysosomes.** COS7 cells were stimulated for 90 min with vehicle (Veh), nigericin (Nig, 10 µM), LLOMe (LL, 1 mM), imiquimod (IQ, 75 µM) or monensin (Mon, 10 µM). **(A)** Immunofluorescence images suggesting that TGN46 can co-localize with LAMP1 after treatment with NLRP3-activating stimuli and monensin. **(B)** Pearson's correlation coefficient (PCC) between TGN46 and LAMP1 for all stimuli. **(C)** Immunofluorescence images showing that stimuli had no effect on co-localization between TGN46 and MICAL-L1. **(D)** Pearson's correlation coefficient (PCC) between TGN46 and MICAL-L1 for all stimuli. Images are representative of 3-5 independent experiments. Blue represents nuclei staining by DAPI. Yellow arrowheads indicate instances of co-localization at the Golgi, white arrowheads indicate co-localization on puncta. Scale bars: 10 µm. Values are mean ± SEM (n = 3). One-way ANOVA followed by Bonferroni's multiple comparison test was performed for data analysis. *: $P \leq 0.05$.

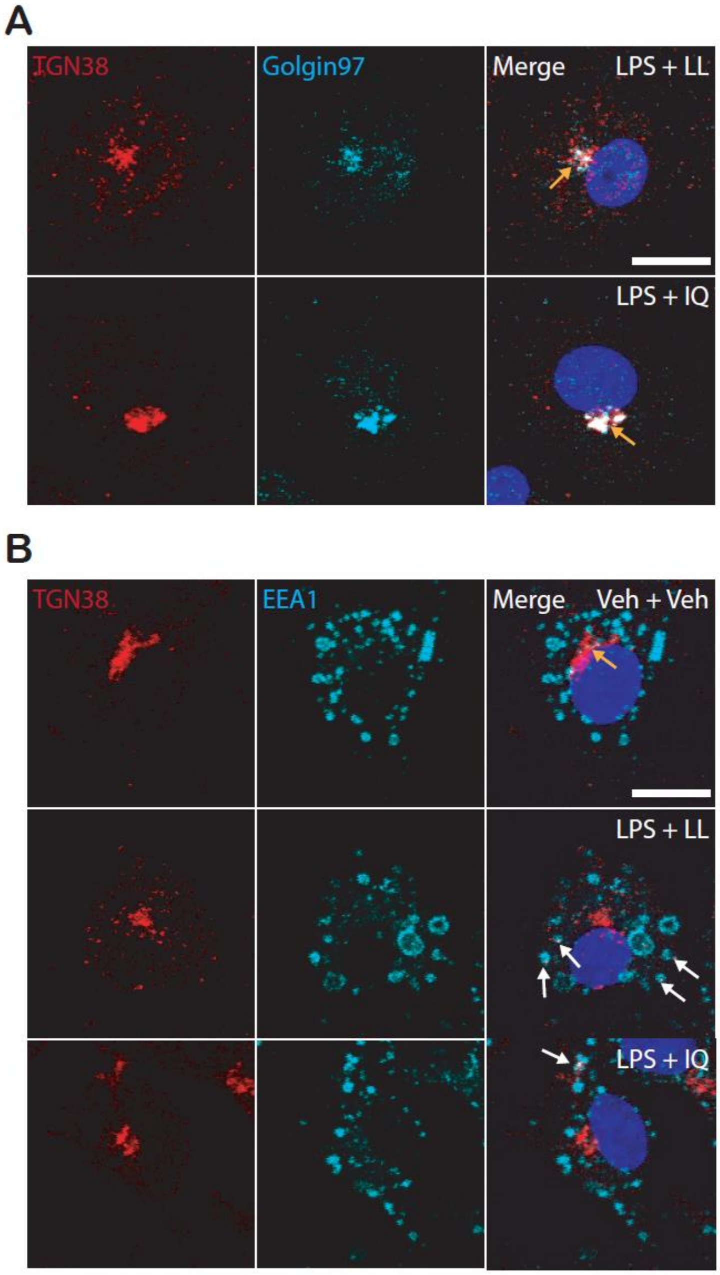


**Fig. S3. TGN38 co-localizes with EEA1 in BMDMs after stimulation with NLRP3-activating stimuli**. BMDMs were primed with LPS (1 µg $mL^{-1}$) for 4 h prior to stimulation. In order to prevent NLRP3-dependent cell death, MCC950 (10 µM) was applied 15 min prior to a 90 min stimulation with LLOMe (LL, 1 mM) or imiquimod (IQ, 75 µM). **(A-B)** Immunofluorescence images showing the co-localization of TGN38 with **(A)** Golgin-97, or **(B)** EEA1. See also Fig 2. Images are representative of 3-5 independent experiments. Blue represents nuclei staining by DAPI. Yellow arrowheads indicate instances of co-localization at the Golgi, white arrowheads indicate co-localization on puncta. Scale bars: 10 µm.

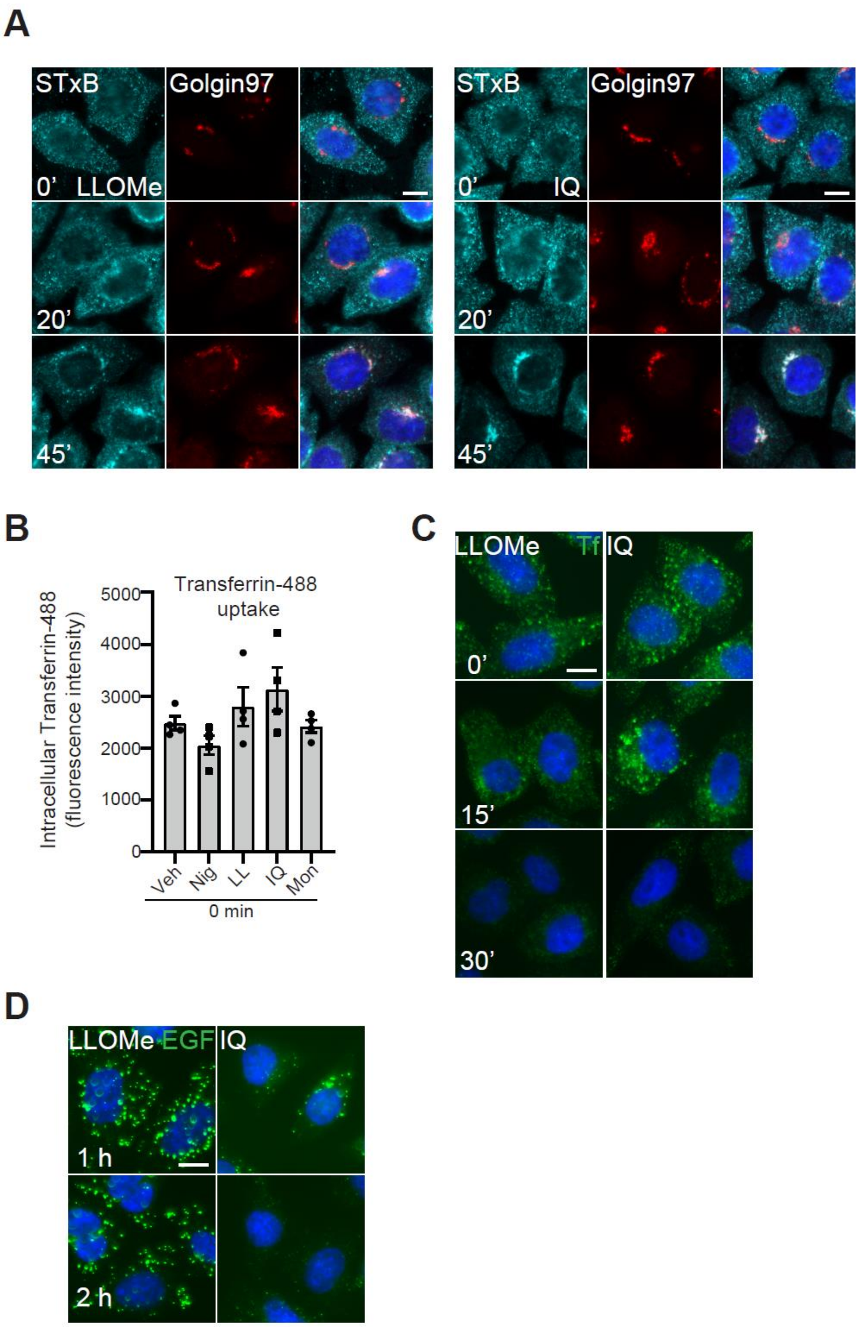


**Fig. S4. Endosomal trafficking is disrupted after NLRP3-activating stimuli. (A)** Representative immunofluorescence images of STxB-Cy3 (cyan) and Golgin-97 (red) in HeLa cells treated with LLOMe (LL, 1 mM) or imiquimod (IQ, 75 µM) from experiments shown in Fig 3B. Quantification of STxB-Cy3 trafficking to the TGN is shown in Fig 3B. **(B)** Transferrin-488 uptake measured in HeLa cells treated with transferrin-488 (5 µg $mL^{-1}$) and with either vehicle (veh), nigericin (Nig, 10 µM), LLOMe (LL, 1 mM), imiquimod (IQ, 75 µM) or monensin (Mon, 10 µM) for 30 min (n = 4). HeLa cells were washed and fixed immediately and transferrin-488 uptake was determined by measuring fluorescence intensity. **(C)** Representative immunofluorescence images of transferrin-488 (Tf, green) in HeLa cells treated with LLOMe or imiquimod from experiments shown in Figure 3E. **(D)** Representative immunofluorescence of

EGF-488 (green) in HeLa cells treated with LLOMe or imiquimod from experiments shown in Fig 3H. Quantification of EGF-488 degradation is shown in Fig 3H. Scale bars: 10 μm. Values are mean ± SEM.

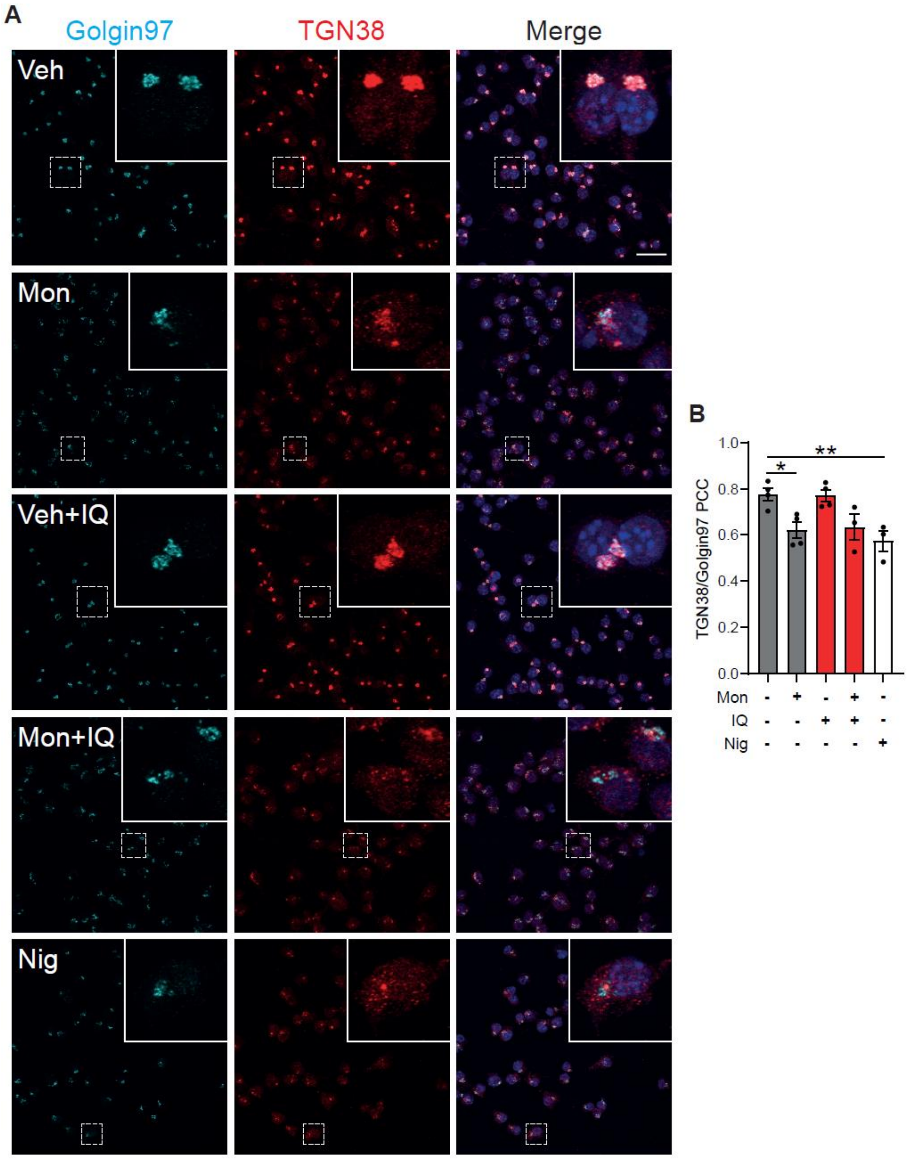


**Fig. S5. Monensin disrupts TGN38 trafficking and TGN structure in iBMDMs.** LPS-primed iBMDMs were treated with vehicle (Veh) or monensin (10 µM) for 1 h prior to MCC950 treatment (10 µM, 15 min) and subsequent imiquimod (IQ, 75 µM) or nigericin (Nig, 10 µM) stimulation for a further 2 h (n = 3-4). **(A)** Immunofluorescence images showing the co-localization of TGN38 with Golgin-97. **(B)** Pearson's correlation coefficient (PCC) between TGN38 and Golgin-97. Blue represents nuclei staining by DAPI. Images were acquired using confocal microscopy. Enlarged images show regions denoted by the white box. Scale bars: 25 µm. Values are mean ± SEM. One-way ANOVA followed by Dunnett's multiple comparison test (versus vehicle) was performed for data analysis. *: P≤ 0.05, **: P≤0.01.

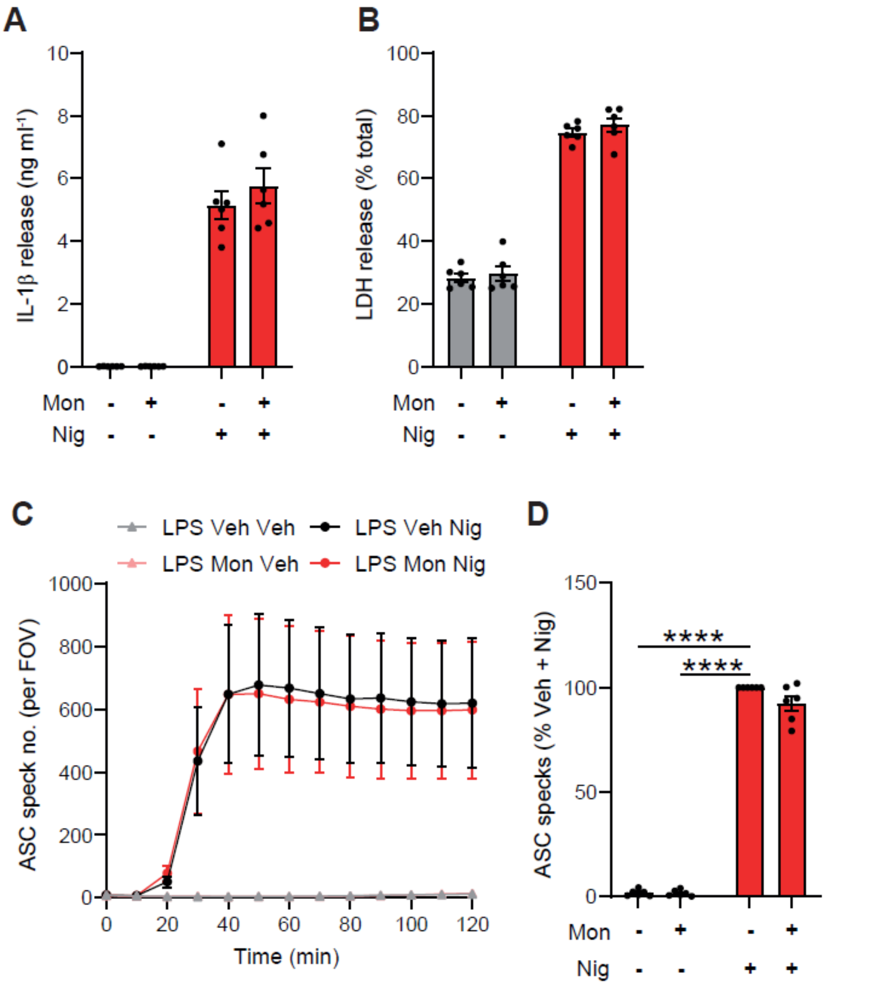


**Fig. S6. Monensin does not potentiate nigericin-induced NLRP3 activation in iBMDMs.** (A-B) LPS-primed iBMDMs were treated with vehicle (Veh) or monensin (Mon, 10 µM) for 1 h prior to nigericin stimulation (Nig, 10 µM) for a further 1 h. Supernatants were assessed for (A) IL-1β and (B) LDH release (n = 6). **(C-D)** LPS-primed iBMDMs stably expressing ASC-mCherry were treated with vehicle or monensin (10 µM) for 2 h prior to Ac-YVAD-CMK (100 µM, 15 min) followed by nigericin stimulation (10 µM) for a further 2 h. (C) ASC speck formation was measured, and (D) ASC speck number at the final time point of 2 h is shown (% of veh + nigericin treatment) (n = 6). IL-1β release was measured by ELISA. Values are mean ± SEM. Unmatched two-way ANOVA followed by Sidak's multiple comparison test (A, B), or one-sample t test versus a value of 100% followed by Holm-Sidak correction (D), were performed for data analysis. ****: P≤0.0001.

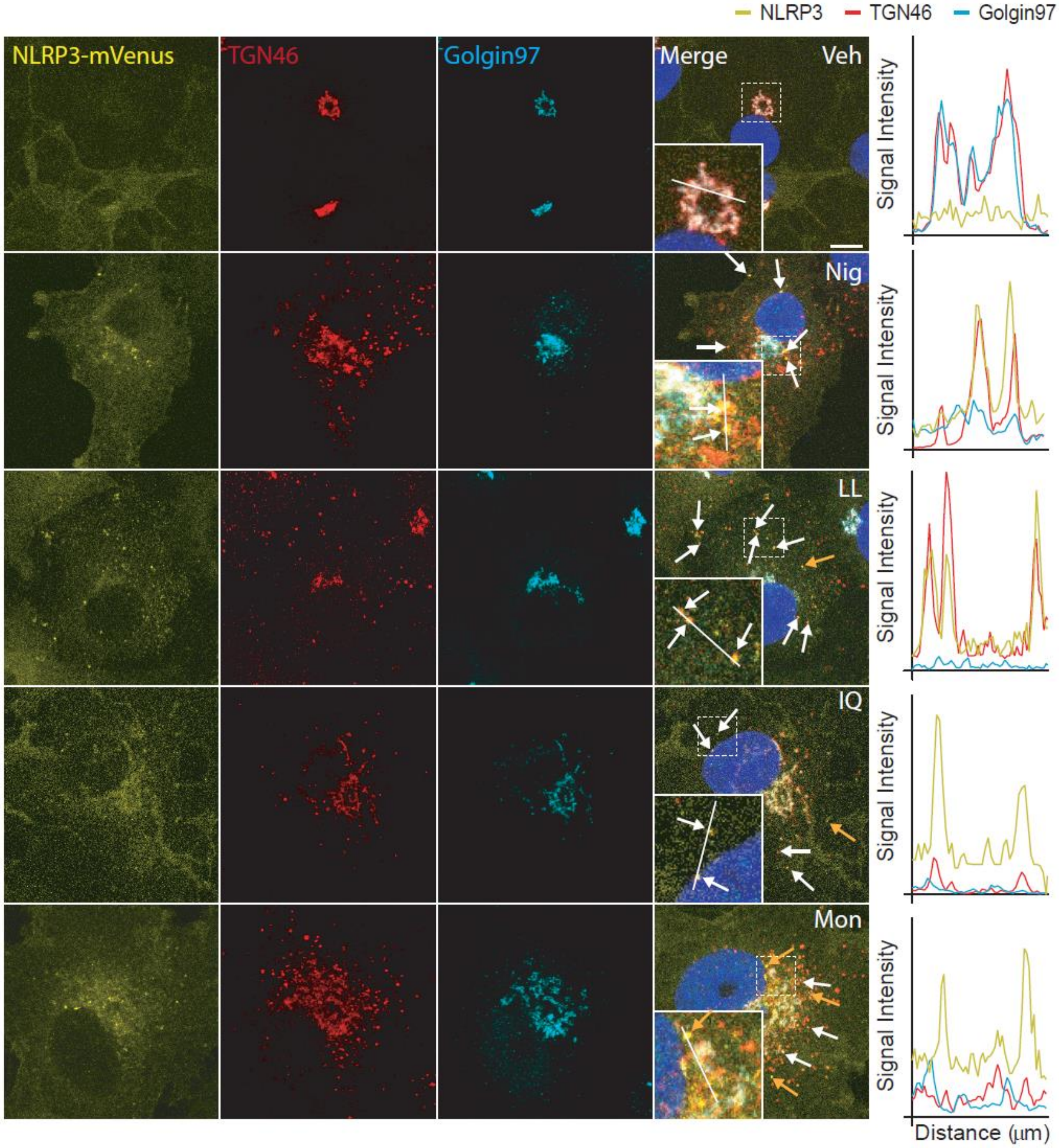


**Fig. S7. NLRP3-mVenus co-localizes with TGN46-positive puncta but not Golgin-97 after stimulation with NLRP3-activating stimuli.** Immunofluorescence images of COS7 cells stably expressing NLRP3-mVenus. Cells were stimulated for 90 min with vehicle (Veh, 0.5% ethanol (v/v)), nigericin (Nig, 10 µM**),** LLOMe (LL, 1mM), imiquimod (IQ, 75 µM**)** or monensin (Mon, 10 µM). Also shown are line graphs depicting changes in fluorescence intensity (min = 0, max = 250) over 10 µm, for NLRP3-mVenus, TGN46 and Golgin97. Images are representative of 3-5 independent experiments. Blue represents nuclei staining by DAPI. White arrow indicates co-localization between NLRP3 and TGN46- positive puncta; yellow arrow indicates NLRP3 puncta alone. Dashed line box highlights area depicted in zoom.

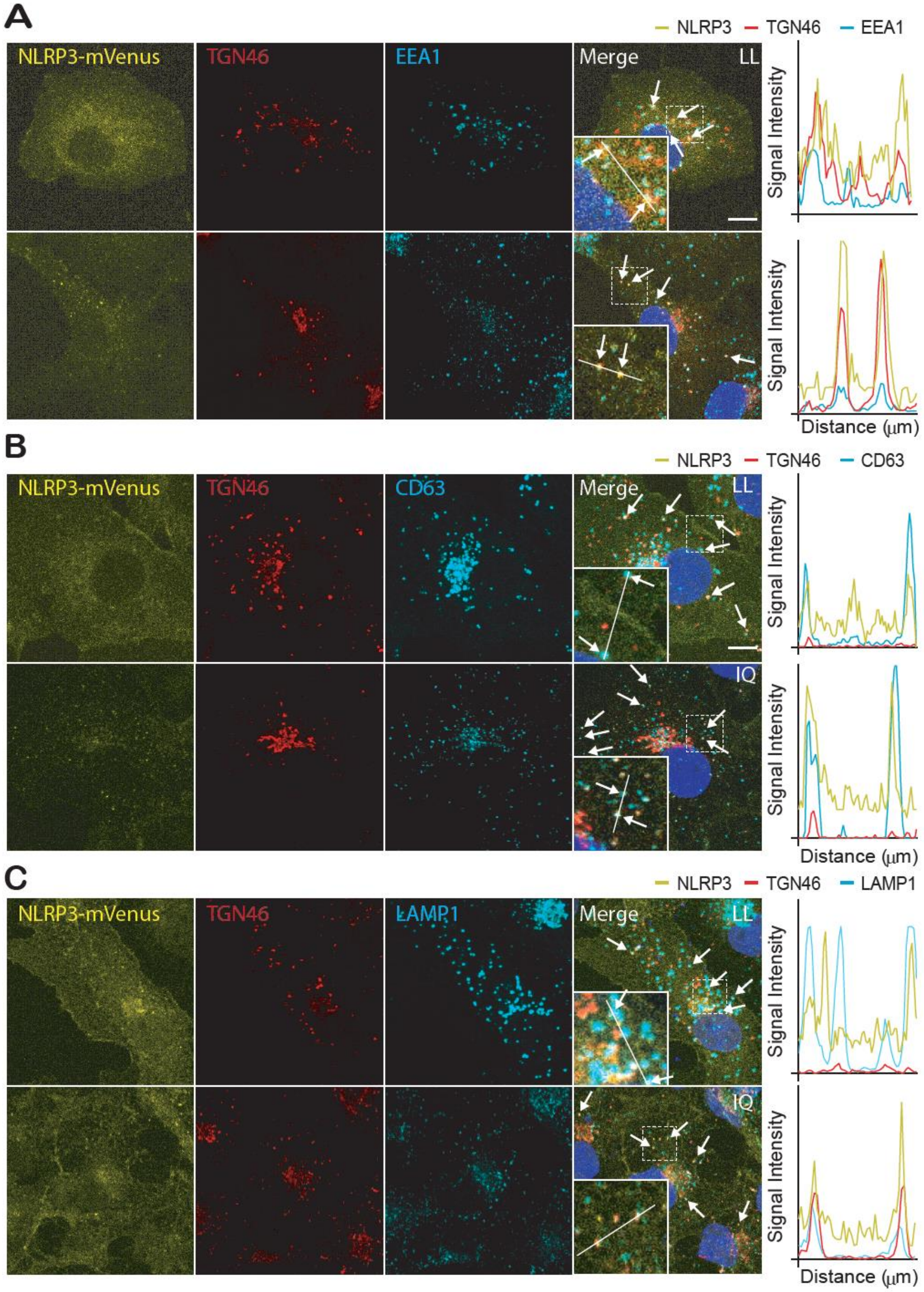


**Fig. S8. NLRP3-mVenus co-localizes with TGN46-, EEA1-, CD63-, LAMP1-positive puncta after stimulation with NLRP3-activating stimuli.** Immunofluorescence images of COS7 cells stably expressing NLRP3-mVenus. Cells were stimulated for 90 min with LLOMe (LL, 1 mM), or imiquimod (IQ, 75 µM)). Also shown are line graphs depicting changes in fluorescence intensity (min = 0, max = 250) over 10 µm, for NLRP3-mVenus, TGN46 and **(A)** EEA1, **(B)** CD63, or **(C)** LAMP1. Images are representative of 3-5 independent experiments. Blue represents nuclei staining by DAPI. White arrow represents co-localization between NLRP3 and **(A)** EEA1-, **(B)** CD63- or **(C)** LAMP1-positive puncta; yellow arrow indicates NLRP3 puncta alone. Dashed line box highlights area depicted in zoom.

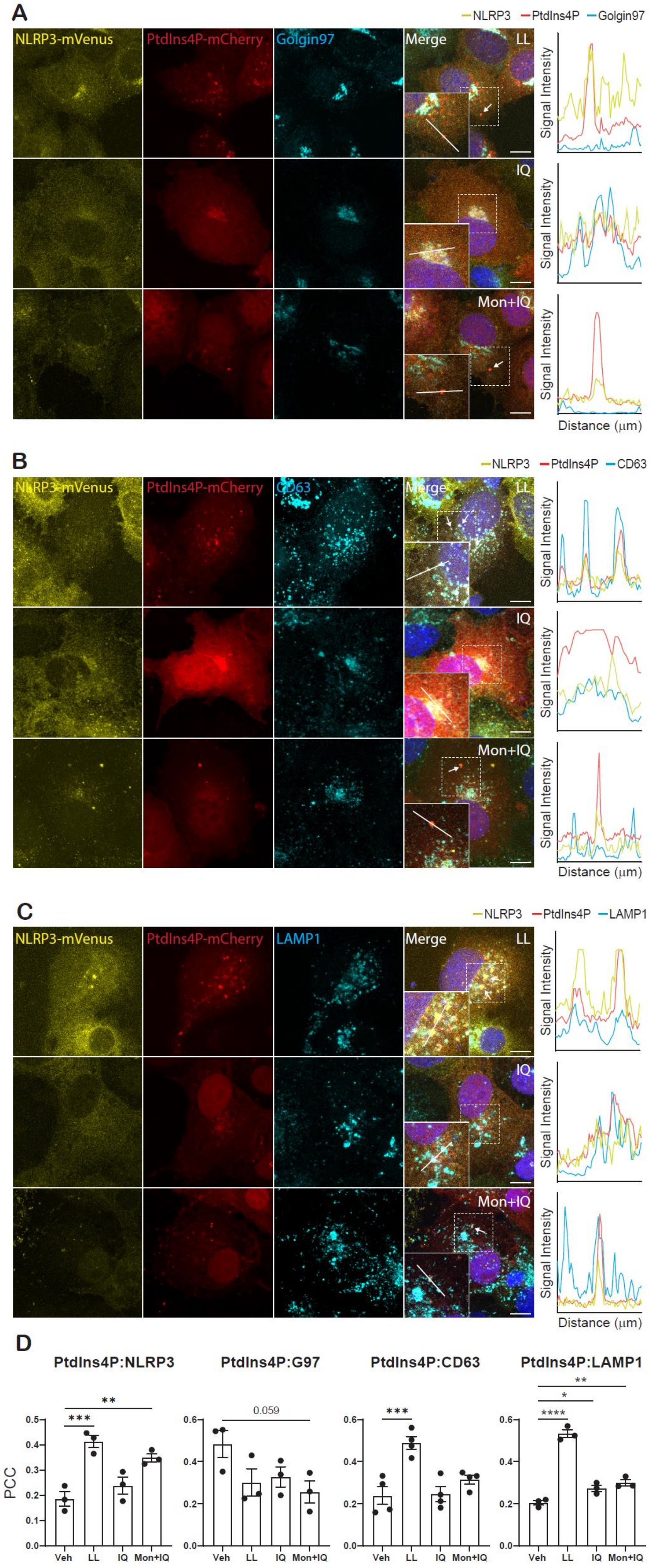

A
NLRP3 PtdIns4P Golgin97
NLRP3-mVenus
PtdIns4P-mCherry
Golgin97
Merge
LL
IQ
Mon+IQ
Signal Intensity
Distance (μm)
B
NLRP3 PtdIns4P CD63
NLRP3-mVenus
PtdIns4P-mCherry
CD63
Merge
LL
IQ
Mon+IQ
Signal Intensity
Distance (μm)
C
NLRP3 PtdIns4P LAMP1
NLRP3-mVenus
PtdIns4P-mCherry
LAMP1
Merge
LL
IQ
Mon+IQ
Signal Intensity
Distance (μm)
D
PtdIns4P:NLRP3
PtdIns4P:G97
PtdIns4P:CD63
PtdIns4P:LAMP1
PCC
0.059
Veh LL IQ Mon+IQ

**Fig. S9. NLRP3-mVenus co-localizes with PtdIns4P on CD63- and LAMP1-positive puncta after stimulation with NLRP3-activating stimuli. (A-C)** Fluorescence and immunofluorescence images of COS7 cells stably expressing NLRP3-mVenus. Cells were stimulated for 90 min with LLOMe (LL, 1 mM), imiquimod (IQ, 75 µM) or monensin (Mon, 10 µM) for 90 min prior to imiquimod stimulation (IQ, 75 µM) for a further 90 min. Co-localization between NLRP3-mVenus, PtdIns4P and **(A)** Golgin-97, **(B)** CD63, or **(C)** LAMP1 was then assessed. Also shown are line graphs depicting changes in fluorescence intensity (min = 0, max = 250) over 10 µm, for each of the conditions and stains. Images are representative of 3-4 independent experiments. Blue represents nuclei staining by DAPI. White arrow indicates co-localization between NLRP3, PtdIns4P, and **(A)** Golgin-97 **(B)** CD63- or **(C)** LAMP1 positive puncta; yellow arrow indicates NLRP3 puncta alone. Dashed line box highlights area depicted in zoom. **(D)** Pearson's correlation coefficient (PCC) between PtdIns4P and the indicated proteins. Values are mean ± SEM, (n=3-4). One-way ANOVA followed by Dunnett's multiple comparison test (versus vehicle) was performed for data analysis. *: $P \leq 0.05$, **: $P \leq 0.01$**,** *** $P \leq 0.001$, **** $P \leq 0.0001$.

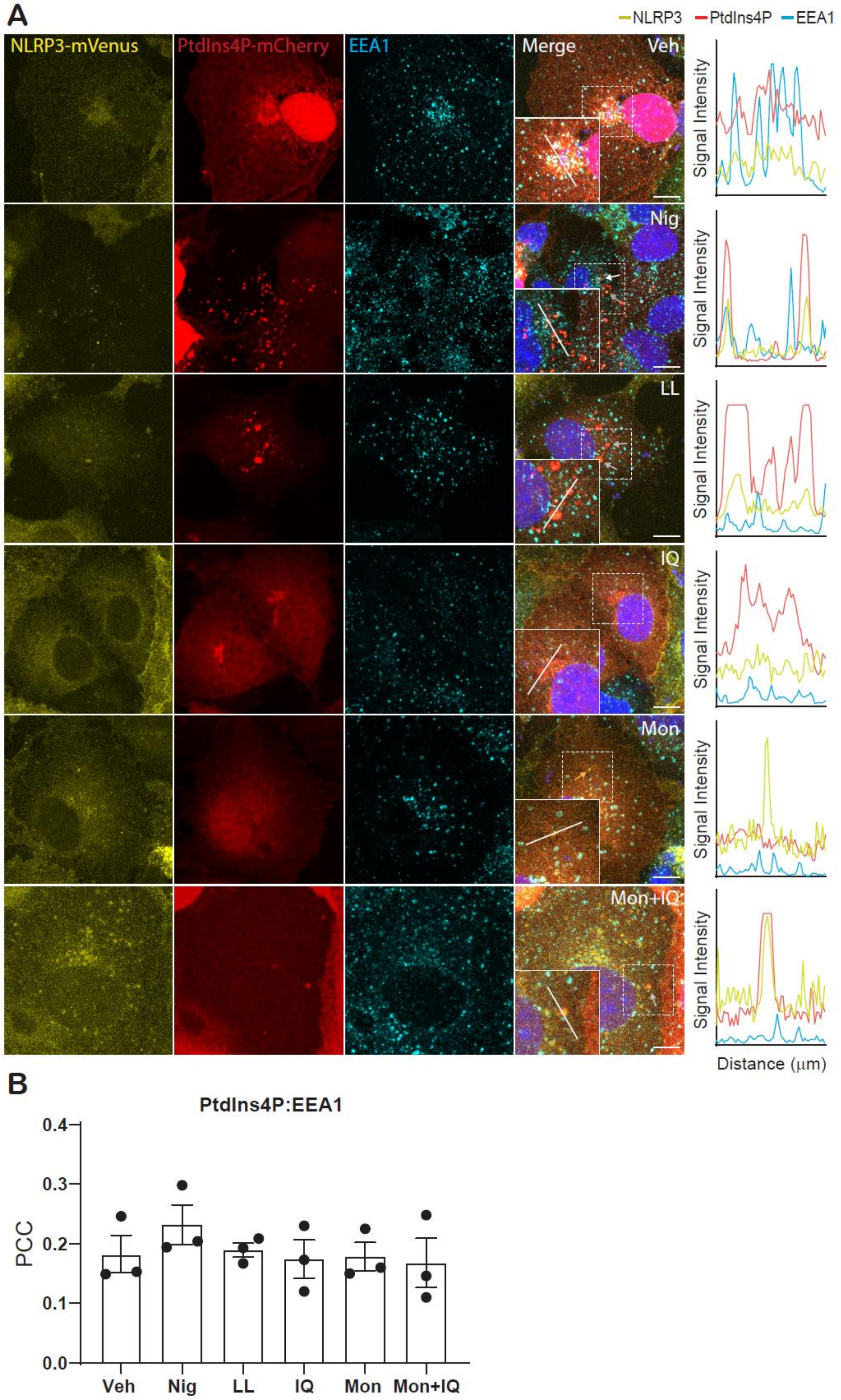
A
NLRP3
PtdIns4P
EEA1
NLRP3-mVenus
PtdIns4P-mCherry
EEA1
Merge
Veh
Nig
LL
IQ
Mon
Mon+IQ
Signal Intensity
Distance (μm)
B
PtdIns4P:EEA1
PCC
0.0
0.1
0.2
0.3
0.4
Veh
Nig
LL
IQ
Mon
Mon+IQ

**Fig. S10. NLRP3 mVenus, PtdIns4P puncta show limited co-localization with EEA1 after stimulation with NLRP3-activating stimuli. (A-C)** Fluorescence and immunofluorescence images of COS7 cells stably expressing NLRP3-mVenus. Cells were stimulated for 90 min with vehicle (Veh, 0.5% ethanol (v/v)), nigericin (Nig, 10 µM), LLOMe (LL, 1 mM), imiquimod (IQ, 75 µM), monensin (Mon, 10 µM), or monensin (Mon, 10 µM) for 90 min prior to imiquimod stimulation (IQ, 75 µM) for a further 90 min. Co-localization between NLRP3-mVenus, PtdIns4P and **(A)** EEA1 was then assessed. Also shown are line graphs depicting changes in fluorescence intensity (min = 0, max = 250) over 10 µm, for each of the conditions and stains. Images are representative of 3 independent experiments. Blue represents nuclei staining by DAPI. White arrow indicates co-localization between NLRP3, PtdIns4P, and **(A)** EEA1 positive puncta; yellow arrow indicates NLRP3 puncta alone; grey arrow indicates co-localization between NLRP3 and PtdIns4P alone. Dashed line box highlights area depicted in zoom. **(B)** Pearson's correlation coefficient (PCC) between PtdIns4P and EEA1. Values are mean ± SEM, (n=3). One-way ANOVA followed by Dunnett's multiple comparison test (versus vehicle) was performed for data analysis**.**
